# BnaA03.WRKY28, interacting with BnaA09.VQ12, acts as a brake factor of activated BnWRKY33-mediated resistance outburst against *Sclerotinia sclerotiorum* in *Brassica napus*

**DOI:** 10.1101/2021.01.28.428601

**Authors:** Ka Zhang, Fei Liu, Zhixin Wang, Chenjian Zhuo, Kaining Hu, Xiaoxia Li, Jing Wen, Bin Yi, Jinxiong Shen, Chaozhi Ma, Tingdong Fu, Jinxing Tu

**Affiliations:** National Key Laboratory of Crop Genetic Improvement, College of Plant Science and Technology, National Sub-Center of Rapeseed Improvement in Wuhan, Huazhong Agricultural University, Wuhan 430070, China; Institute of Vegetable, Wuhan Academy of Agricultural Sciences, Wuhan 430065, China

## Abstract

*Sclerotinia sclerotiorum* causes substantial damage to the growth of *Brassica napus* (rapeseed) and makes a significant loss of crop yield. The plant innate immune system may be the primary solution to defense against *S. sclerotiorum* for rapeseed. Here, we identify that BnWRKY33, a transcription factor in the innate immune pathway, can be rapidly phosphorylated and activated by the MAPK cascade after rapeseed is infected with *S. sclerotiorum*. In the MAPK cascade, activated BnaA03.MKK4 phosphorylates and activates BnaA06.MPK3 and BnaC03.MPK3. The activated BnMPK3 acts on the substrate BnWRKY33 to enhance its transcriptional activity and trigger a transcriptional burst of *BnWRKY33*, which helps plants effectively resist the pathogenic fungi by enhancing the expression of phytoalexin synthesis-related genes. With constant infection, *BnaA03.WRKY28* and *BnaA09.VQ12* are induced, and BnaA03.WRKY28 physically interacts with BnaA09.VQ12 to form a protein complex. BnaA03.WRKY28 preferentially binds to the promoter of *BnWRKY33* with the help of BnaA09.VQ12. Compared with activated BnWRKY33, BnaA03.WRKY28 has a lower transcriptional activity on downstream *BnWRKY33*, which leads to weaker resistance against *S. sclerotiorum* for rapeseed in the later stage of infection. Furthermore, the induced *BnaA03.WRKY28* may promote axillary bud activity and axillary meristem initiation by regulating the expression of branching-related genes (such as *BnBRC1*), thus promoting the formation of branches in the leaf axils.

**One-sentence summary:** Under constant infection by *Sclerotinia sclerotiorum*, BnaA03.WRKY28 interacts with BnaA09.VQ12 and takes precedence over phosphorylated BnWRKY33 to bind to the *BnWRKY33* promoter, thereby weakening resistance but promoting branching.

## INTRODUCTION

In nature, plants are frequently attacked by pathogens. When challenged by pathogens, the innate immune system of plants is activated (Dangl and Jones, 2001; Jones and Dangl, 2006). The plant innate immune system has evolved into two branches, pathogen-associated molecular pattern (PAMP)-triggered immunity (PTI) and effector-triggered immunity (ETI) (Jones and Dangl, 2006; Tsuda and Katagiri, 2010).

*Brassica napus* (rapeseed) is one of the most important oil crops worldwide. However, *Sclerotinia sclerotiorum* is generally considered one of the most economically damaging diseases on rapeseed (Kharbanda and Tewari, 1996; Derbyshire and Denton-Giles, 2016). *S. sclerotiorum* is a typical necrotrophic fungal pathogen with a wide range of hosts, including many important economic crops such as rapeseed, peanut, soybean, and sunflower (Bolton et al., 2006). Great efforts have been made to identify quantitative genetic loci (QTL) defense against *S. sclerotiorum* in *Brassica napus* (Zhao et al., 2006; Wei et al., 2016; Zhang et al., 2019). However, the lack of germplasms immune to Sclerotinia in *Brassica napus* and all Brassicaceae species makes the identification of QTLs related to Sclerotinia resistance and the breeding of disease-resistant varieties less effective, and disease control in *Brassica napus* primarily relies on partially resistant lines (Wang et al., 2018; Zhang et al., 2019). Transgenic manipulation of genes associated with the innate immune system provides alternative approaches for a deeper understanding of rapeseed response to *S. sclerotiorum* and the production of rapeseed varieties with strong resistance to this fungal pathogen.

Plants respond to the invasion of pathogens by regulating the expression of numerous genes, in which transcription factors (TFs) are widely involved (Singh et al., 2002). The WRKY superfamily TFs, which contain conserved WRKYGQK together with an atypical zinc-finger structure, respond to diverse abiotic and biotic stresses with the ability to specifically bind to T/CTGACC/T (W-box) *cis*-acting elements in the promoter of target genes (Eulgem et al., 2000; Ülker and Somssich, 2004; Rushton et al., 2010). In *Arabidopsis thaliana*, WRKY18, WRKY40, and WRKY60 are involved in seed germination through ABA signaling (Shang et al., 2010); WRKY46, WRKY54, and WRKY70 function as negative regulators of BR-regulated drought tolerance (Chen et al., 2017). Additionally, *AtWRKY33* plays a critical role in widespread stress responses. The *wrky33* mutant is sensitive to NaCl stress (Jiang and Deyholos, 2009), and WRKY33 can bind to *RAP2.2* to regulate its expression during submergence induced hypoxia response (Tang et al., 2020). Overexpression of *AtWRKY33* enhances plants resistance against the necrotrophic fungal pathogen *Botrytis cinerea* (Birkenbihl et al., 2012; Liu et al., 2017). WRKY33 regulates the expression of *CYP71B15* (*PAD3*) and *CYP71A13*, which encode cytochrome P450 enzymes required for the synthesis of the antimicrobial phytoalexin camalexin, and WRKY33 can directly target the W-box in the promoter of *PAD3* (Qiu et al., 2008; Mao et al., 2011). Moreover, WRKY33 regulates its own expression by binding to *WRKY33* promoter (Mao et al., 2011). In *Brassica napus*, *BnWRKY33* has been reported to play a positive regulatory role in Sclerotinia resistance (Wang et al., 2014b; Liu et al., 2018). *BnWRKY33* overexpression lines display more resistant to *S. sclerotiorum* than wild-type plants, and as in Arabidopsis, BnWRKY33 can bind to its own promoter. Nonetheless, our knowledge about the mechanism by which *BnWRKY33* positively defends against *S. sclerotiorum* is limited.

Both PTI and ETI can trigger the activation of kinase cascades mediated by the mitogen-activated protein kinase (MAPK) class (Tsuda and Katagiri, 2010; Bigeard et al., 2015; Cui et al., 2015). Plant MAPK cascade-mediated signaling is an essential event in response to pathogens. The basic MAPK cascade consists of three graded kinases. MAPKs are phosphorylated and activated by their upstream MAPK kinases (MAPKKs/MKKs), and MAPKKs are activated by their upstream MAPKK kinases (MAPKKKs) through the phosphorylation of conserved Ser/Thr in the activation loop ((S/T)XXXXX(S/T)) of plant MAPKKs (Doczi et al., 2012; Meng and Zhang, 2013). The MKK4/MKK5-MPK3/MPK6-mediated MAPK signaling cascade plays an essential role in plant immune response (Asai et al., 2002; Zhang et al., 2018). Furthermore, in Arabidopsis, activated MPK3/6 can directly phosphorylate downstream substrates, such as WRKY33, ERF104, and VIP1, to induce defense-related genes to respond to pathogens (Djamei et al., 2007; Bethke et al., 2009; Mao et al., 2011).

WRKY superfamily TFs not only play roles in the pathogen-induced defense response, but are also involved in plant growth and development (Eulgem et al., 2000; Rushton et al., 2010). For example, *AtWRKY75* is associated with lateral root length and root hair number (Devaiah et al., 2007). Overexpression of *AtWRKY71* significantly enhanced the activity of axillary meristem and produced more branches (Guo et al., 2015). Additionally, *WRKY28* is reported to involve in the fate determination of megaspore mother cells (MMCs) and the regulation of leaf senescence in Arabidopsis. In the development process that restricts the differentiation of multiple adjacent somatic cells of MMC into MMCs, *AtWRKY28* is specifically expressed in the hypodermal somatic cells surrounding the MMC, inhibiting these somatic cells from gaining the fate of MMC-like fate. The cytochrome P450 gene *KLU* promotes *WRKY28* expression through chromatin remodeling complex SWR1-mediated H2A.Z deposition at *WRKY28* (Zhao et al., 2018). Besides, *WRKY28* positively regulates salicylic acid (SA) biosynthesis by promoting the expression of SA biosynthesis-related genes (van Verk et al., 2011). A light-signaling protein, FAR-RED ELONGATED HYPOCOTYL3 (FHY3), can physically bind to the promoter region of *WRKY28* and inhibit the expression of *WRKY28*, thereby negatively regulating SA biosynthesis and leaf senescence (Tian et al., 2020). In contrast, *BnWRKY28* (orthologous gene of *AtWRKY28*) is reported to be induced by *S. sclerotiorum* and JA/ETH in *Brassica napu*s (Yang et al., 2009). However, the specific biological functions and molecular mechanisms of BnWRKY28 remain elusive.

VQ proteins, which contain a conserved FxxxVQxLTG motif, are involved in developmental processes (Li et al., 2014; Lei et al., 2017), biotic stress responses (Wang et al., 2015; Chen et al., 2018b), and abiotic stress responses (Perruc et al., 2004; Hu et al., 2013). AtVQ12 was identified to negatively regulate the resistance to *Botrytis cinerea* in Arabidopsis (Wang et al., 2015). AtVQ12 overexpression plants show susceptible to *B. cinerea*. A common feature of most VQ proteins is that they interact with WRKY TFs as co-factors (Cheng et al., 2012; Jing and Lin, 2015). For example, VQ9 and WRKY8 form a complex that contributes to salinity tolerance (Hu et al., 2013). On the other hand, VQ10 interacts with WRKY8 and participates in plant defense against the fungal pathogen *B. cinerea* (Chen et al., 2018b).

In this study, we demonstrate that one copy of *BnWRKY28* (*BnaA03.WRKY28*) plays a negative role in defense against *S. sclerotiorum* in *Brassica napu*s, and BnaA03.WRKY28 directly binds to the promoter of *BnWRKY33*. BnWRKY33 is a substrate of BnMPK3, and BnWRKY33 can be phosphorylated and activated downstream of the MAPK cascade when rapeseed is infected with *S. sclerotiorum.* In addition, BnaA03.WRKY28 and BnWRKY33 competitively bind to the promoter of *BnWRKY33* in the late stage of the Sclerotinia response. However, the formation of a protein complex between BnaA03.WRKY28 and BnaA09.VQ12 enhances the binding affinity of BnaA03.WRKY28 to the *BnWRKY33* promoter, which helps BnaA03.WRKY28 act as a brake factor in resistance to *S. sclerotiorum* in *Brassica napus*.

## RESULTS

### BnaA03.WRKY28 Negatively Regulates the Resistance against Sclerotinia sclerotiorum in Brassica napus

Many BnWRKY transcription factors (TFs) respond to *Sclerotinia sclerotiorum* and hormone treatments in *Brassica napus*, including *BnWRKY28,* which was highly induced after the inoculation of *S. sclerotiorum* (Yang et al., 2009). *Brassica napus* is an allopolyploid, including the AA and CC subgenomes (Chalhoub et al., 2014). Using the homologous cloning method, we identified five copies of *BnWRKY28* on chromosomes A01, C01, A03, A08, C08 in Jia9709 and Westar, named *BnaA01.WRKY28*, *BnaC01.WRKY28*, *BnaA03.WRKY28*, *BnaA08.WRKY28*, and *BnaC08.WRKY28*, respectively, all of which are orthologous to *AtWRKY28* based on phylogenetic analysis (Supplemental Figure 1A, 1B). To investigate whether *BnWRKY28* plays a role in response to *S. sclerotiorum* in *Brassica napus*, we performed quantitative (q) RT-PCR analysis of transcripts abundance when wild-type (WT) plants were challenged with this fungal pathogen. We found that of all the copies, only the A03 copy was strongly and dramatically induced (Figure 1A). *BnaA03.WRKY28* was gradually induced with the inoculation time, and the expression level peaked at 48 hours (h) after inoculation, followed by a decreased expression trend. Salicylic acid (SA) and jasmonic acid (JA) are two essential defense hormones (Pieterse et al., 2012; Shigenaga et al., 2017). *BnaA03.WRKY28* was also induced when treated with SA and JA, but it peaked at 24 h by 50 µM MeJA and 12 h by 1 mM SA (Figure 1B). These data indicate that *BnaA03.WRKY28* respond to *S. sclerotiorum* and may play a role in plants suffering from infection by this pathogen.

**Figure 1.**
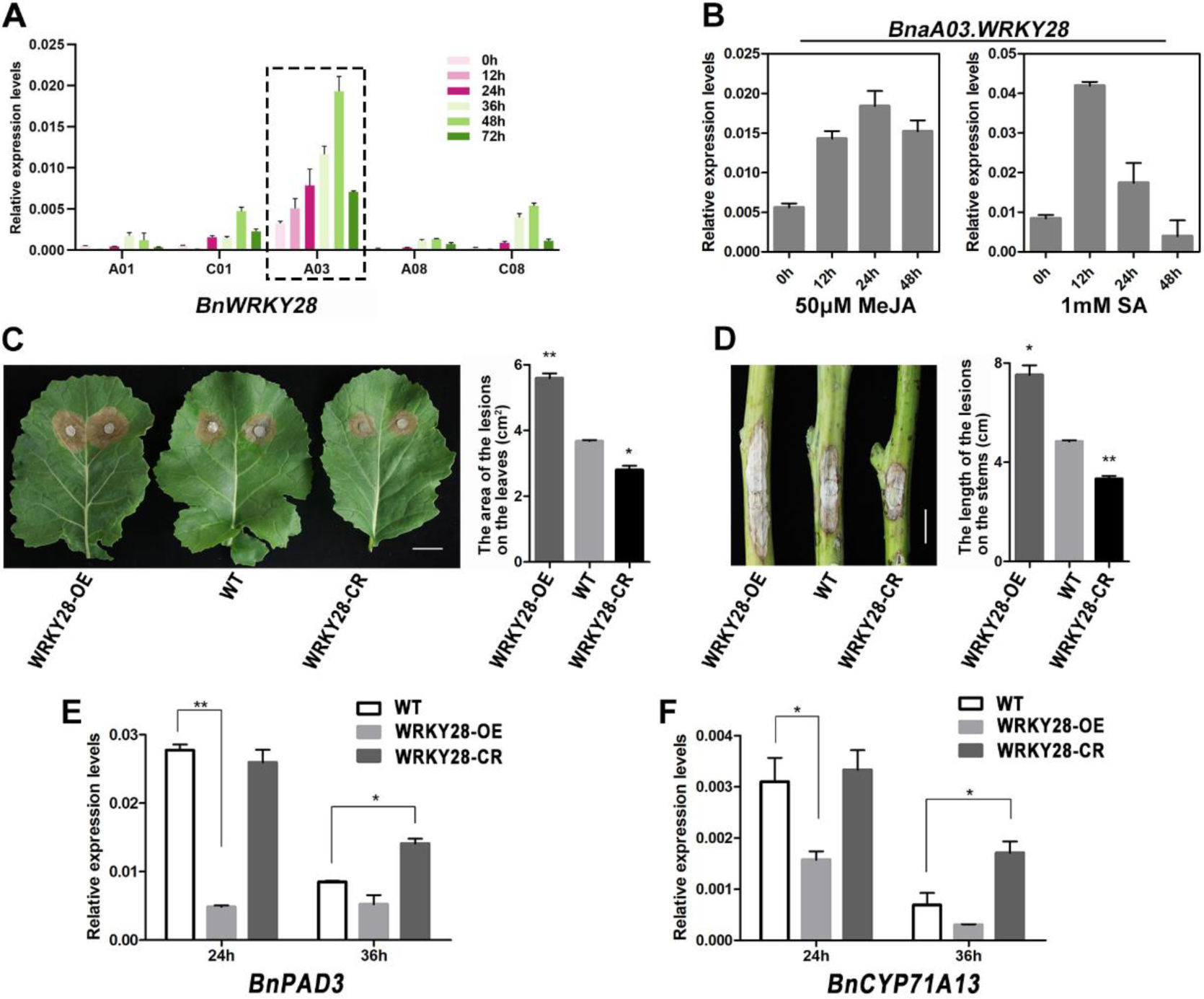
Negative Effect of *BnaA03.WRKY28* on *S. sclerotiorum* Resistance in *Brassica napus*. **(A)** Induced expression profiles of five copies (A01, C01, A03, A08, and C08) of *BnWRKY28* were identified at 0 h, 12 h, 24 h, 36 h, 48 h, and 72 h after inoculation with *S. sclerotiorum*. The latest fully unfolded leaves of rapeseed (Jia 9709) were used for infection for approximately 4 weeks. Data are shown as means ± SD (*n* = 3). **(B)** The changes of transcript abundance of *BnaA03.WRKY28* were tested when rapeseed plants (approximately 4 weeks) were treated with 50 µM MeJA or 1 mM SA after 0 h, 12 h, 24 h, 36 h, and 48 h. Data are shown as means ± SD (*n* = 3). **(C)** The lesions of detached leaves from *BnaA03.WRKY28* transgenic T3 homozygous lines and wild-type (WT) plants (Jia 9709) at approximately 4 weeks were imaged and lesion areas were calculated when inoculated with 7 mm *S. sclerotiorum* hyphae agar block at 22°C for 48 h. Scale bar, 2 cm. **(D)** The lesion lengths on the stems of transgenic lines and WT at the flowering stage were imaged and calculated when inoculated with *S. sclerotiorum* for 4 days. Scale bar, 2 cm. **(E)** and **(F)** Relative expression levels of two camalexin biosynthesis-related genes, *BnPAD3* and *BnCYP71A13*, in *BnaA03.WRKY28* transgenic lines and WT plants after *S. sclerotiorum* inoculation for 24 h or 36 h. In **(C)** to **(F)**, WRKY28-OE indicated *BnaA03.WRKY28* overexpression lines, and WRKY28-CR indicated *BnaA03.WRKY28* editing lines mediated by the CRISPR/Cas9 system. Data are shown as means ± SD (*n* = 3). Asterisks indicate significant differences compared with WT (t-test, \**P* < 0.05, \*\**P* < 0.01).

To further elucidate the biological function of *BnaA03.WRKY28* in the *S. sclerotiorum* response, we generated *BnaA03.WRKY28* overexpression (WRKY28-OE) lines and CRISPR/Cas9 mediated *BnWRKY28* editing (WRKY28-CR) lines. We examined the expression levels of *BnaA03.WRKY28* in overexpressing lines and the edited form of *BnaA03.WRKY28* in CRISPR/Cas9 lines, and measured the lesion area of overexpression lines and homozygous editing lines compared with WT at 48 h after inoculation with *S. sclerotiorum* (Supplemental Figure 2C to 2K). The WRKY28-OE plants showed more severe disease symptoms and fungal invasion than WT during infection, whereas the WRKY28-CR plants exhibited enhanced resistance compared with WT (Figure 1C and 1D, Supplemental Figure 2A and 2B). We also analyzed the pathogen-induced expression of camalexin biosynthetic genes, including *BnPAD3* and *BnCYP71A13* (Figure 1E and 1F). At 24 h after infection with *S. sclerotiorum*, the transcripts of *BnPAD3* and *BnCYP71A13* in WRKY28-CR lines were similar to that of WT, but significantly reduced in WRKY28-OE lines. In contrast, at 36 h after infection, the expression of *BnPAD3* and *BnCYP71A13* in WRKY28-CR lines remained at a relatively high level compared with WT. The differences of transcripts accumulation of these genes indicated that the synthesis of camalexin, an important phytoalexin, may be reduced in WRKY28-OE lines compared with WT, but increased in WRKY28-CR lines. Taken together, these data suggest that *BnaA03.WRKY28* may be a negative regulator of Sclerotinia resistance in *Brassica napu*s.

### BnaA03.WRKY28 Directly Targets *BnWRKY33*

To explore the potential mechanism by which *BnaA03.WRKY28* plays a negative regulatory role in Sclerotinia resistance, we performed RNA-seq and chromatin immunoprecipitation sequencing (ChIP-seq) analyses using *BnaA03.WRKY28* fused FLAG tag overexpression lines. The fused target protein can be stably expressed in line Z95 (Supplemental Figure 3A). In ChIP-seq, approximately 47,000 *BnaA03.WRKY28*-bound peaks were identified (Supplemental Table 1). Most of the binding sites of these peaks were located at promoter regions (Supplemental Figure 3B), with strong enrichment of the TTGACT/C motif (Supplemental Figure 3C). In RNA-seq, 3,972 up-regulated genes and 3,322 down-regulated genes were identified (Supplemental Table 2). Gene ontology (GO) analysis of up-regulated and down-regulated gene clusters indicated that differentially expressed genes (DEGs) were significantly enriched in response to stimulus/stress (Supplemental Figure 3D and 3E). Of the 7,294 DEGs identified by RNA-seq, the promoters of 3,980 genes were shown to be BnaA03.WRKY28-bound in the ChIP-seq assay (Supplemental Figure 3F; Supplemental Table 3). Among the 3980 overlapping genes, most genes were involved in stress response (Supplemental Figure 3G), and there were about 360 transcription factors encoding genes (Supplemental Figure 3H, Supplemental Table 3). Peaks were found in the promoters of four *BnWRKY33* copies included in the DEGs of RNA-seq (Figure 2A).

**Figure 2.**
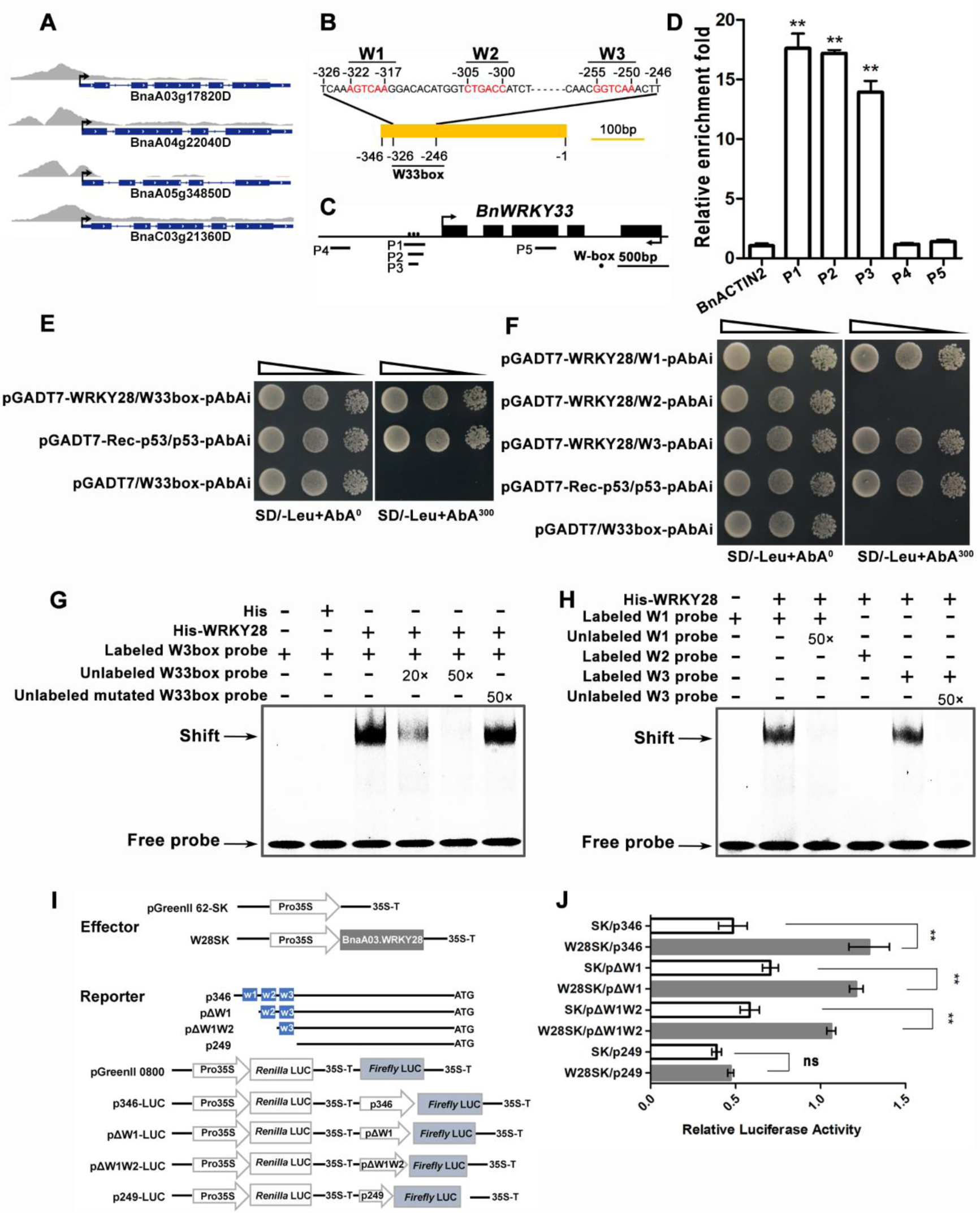
BnaA03.WRKY28 Binds to the Promoter of *BnWRKY33 in vivo* and *in vitro*. **(A)** ChIP-seq showing that the promoter regions of four copies of *BnWRKY33* that were specifically enriched in the DNA immunoprecipitated by BnaA03.WRKY28. The Z95 line (*BnaA03.WRKY28* fusion FLAG tag driven by CaMV35S) was used to perform ChIP-seq. **(B)** Characterization of the *BnWRKY33* promoter sequence. The core sequence of the W-box is marked by red. **(C)** A diagram of the *BnWRKY33* locus showing the positions of amplicons (P1- P5) used for ChIP-qPCR. W-box elements are marked with circles. **(D)** ChIP-qPCR shows that BnaA03.WRKY28 directly binds to the W33box region of *BnWRKY33*. *BnACTIN2*, the promoter region distant from the start codon (P4), and the exon region of *BnWRKY33* (P5) were used as negative controls. Asterisks indicate significant differences compared with *BnACTIN2* (t- test, \*\**P* < 0.01). **(E)** and **(F)** Yeast one-hybrid assays to explore the binding capacity of BnaA03.WRKY28 to *BnWRKY33* promoter. DNA fragments from *BnWRKY33* promoter were cloned into the pAbAi vector, and then transformed into the Y1HGold yeast strain. The coding sequence of BnaA03.WRKY28 was cloned into the pGADT7 vector (pGADT7-WRKY28). Yeast cells cotransformed with pGADT7-BnaWRKY28 and W-box-pAbAi were cultured on selective dropout medium without leucine (SD/-Leu) containing 0 or 300 ng/mL AbA. Leu, leucine; AbA^300^ indicates that the medium contains 300 ng/mL AbA, whereas AbA^0^ indicates that the medium does not contain AbA. The p53 was used as a positive control. The oblique triangles above the images indicate 10^0^, 10^-1^, and 10^-2^ concentration gradient of Yeast cells. **(G)** and **(H)** Electrophoretic mobility shift assays (EMSA) to test the interaction between BnaA03.WRKY28 and the *BnWRKY33* promoter *in vitro*. Probes labeled with Cy5 were used. The mutated probe replaced the core W-box sequence (W1, W2, and W3) with GAGAGA. **(I)** Schematic diagram of effector and reporter constructs using the pGreenII vector set. The coding sequence of transcription factors were introduced into the pGreenII 62-SK (SK) vector as effectors. Truncated promoter regions of *BnWRKY33* were inserted in front of the *firefly luciferase* (LUC) gene as reporters. *Renilla* LUC was used as internal control. **(J)** Dual luciferase assays were conducted to detect the relative luciferase activity (the ratio of firefly luciferase activity to *Renilla* luciferase activity) when cotransformed with effector and reporter. Cotransformation of empty pGreenII 62-SK plasmids and reporter constructs were used as negative controls. Data are shown as means ± SD (*n* = 3). Asterisks indicate significant differences compared with the negative controls (t-test, \*\**P* < 0.01).

WRKY TFs can bind to W-box *cis*-elements with the core sequence 5’- T/CTGACC/T-3’ (Eulgem et al., 2000). *BnWRKY33* promoter contains three W-box elements in the -346 bp to -1 bp region (Figure 2B), named W1, W2, and W3, respectively. Additionally, a region from -326 bp to -246 bp containing three W-boxes was defined as W33box. To confirm whether *BnaA03.WRKY28* binds to the promoter of *BnWRKY33 in vivo*, ChIP-qPCR experiments were performed. We conducted qPCR using primers corresponding to five regions of *BnWRKY33* (Figure 2C) and *BnACTIN2* as a negative control. Amplicons P1 (-370 bp to -166 bp), P2 (-330 bp to -181 bp), P3 (-328 bp to -233 bp), all covering the W33-box, were significantly enriched compared to either amplicon P4 (-1085 bp to -894 bp, distant from W33-box), or amplicon P5 (located in gene body), or the negative control *BnACTIN2* (Figure 2D). In the yeast one-hybrid (Y1H) assay, the bait strain pW33-box-AbAi was transformed with the prey construct pGADT7-Rec-BnaA03.WRKY28 and could grow on SD/-Leu medium containing 300 ng/mL Aureobasidin A (AbA) (Figure 2E), indicating that *BnaA03.WRKY28* can bind to the W33-box in the *BnWRKY33* promoter region. Moreover, by truncating and mutating the W33-box, it was found that W1 and W3, but not W2, were the target sites of BnaA03.WRKY28 (Figure 2F, Supplemental Figure 4A and 4B).

To further confirm that BnaA03.WRKY28 could directly target the cis-acting elements of *BnWRKY33*, electrophoretic mobility shift assays (EMSA) were performed. When we added BnWRKY28 protein, a specific shifted band was produced, and the shifted band was completely competitive when the unlabeled cold probe was added (Figure 2H and 2H). We also detected the effect of BnaA03.WRKY28 on *BnWRKY33* transcription using dual-luciferase transient transcriptional activity assays. The *Firefly* luciferase (LUC) reporter gene was activated when *Pro35S:BnaA03.WRKY28* was co-introduced into Arabidopsis protoplasts with *p346:LUC*, *pΔW1:LUC*, or *pΔW1W2:LUC* (Figure 2I and 2J). These results suggest that BnaA03.WRKY28 can directly target W1 and W3 in the promoter region of *BnWRKY33* and activate the expression of the transcription factor *BnWRKY33*.

### Phosphorylated Activation of BnWRKY33 is Necessary to Enhance Resistance to *S. sclerotiorum* in *Brassica napus*

WRKY33 positively regulates pathogen resistance by binding to its own promoter to upregulate its expression after Arabidopsis was infected with *B. cinerea,* and rapeseed was infected with *S. sclerotiorum* (Mao et al., 2011; Liu et al., 2018). Interestingly, we demonstrated that BnaA03.WRKY28 could also bind to the promoter of *BnWRKY33* and activated the expression of *BnWRKY33*. Moreover, in dual-luciferase transient transcriptional activity assays, when W28SK and W33SK were cotransformed into Arabidopsis protoplasts with p346, it was determined that the relative LUC activity of W28SK/p346 was slightly higher than that of W33SK/p346, suggesting that BnaA03.WRKY28 may have a stronger activation effect on *BnWRKY33* than BnWRKY33 (Figure 3A). We used another set of vectors to detect the transcriptional activity of BnaA03.WRKY28 and BnWRKY33 (Figure 3B). It was revealed that the transcriptional activity of BnaA03.WRKY28 was not lower than that of BnWRKY33 (Figure 3C). The expression level of *BnWRKY33* in *BnaA03.WRKY28* overexpression lines were higher than that in WT (Figure 3D). It was indicated that BnaA03.WRKY28 should be a positive factor in Sclerotinia resistance, similar to *BnWRKY33*, which appeared to contradict our aforementioned description.

**Figure 3.**
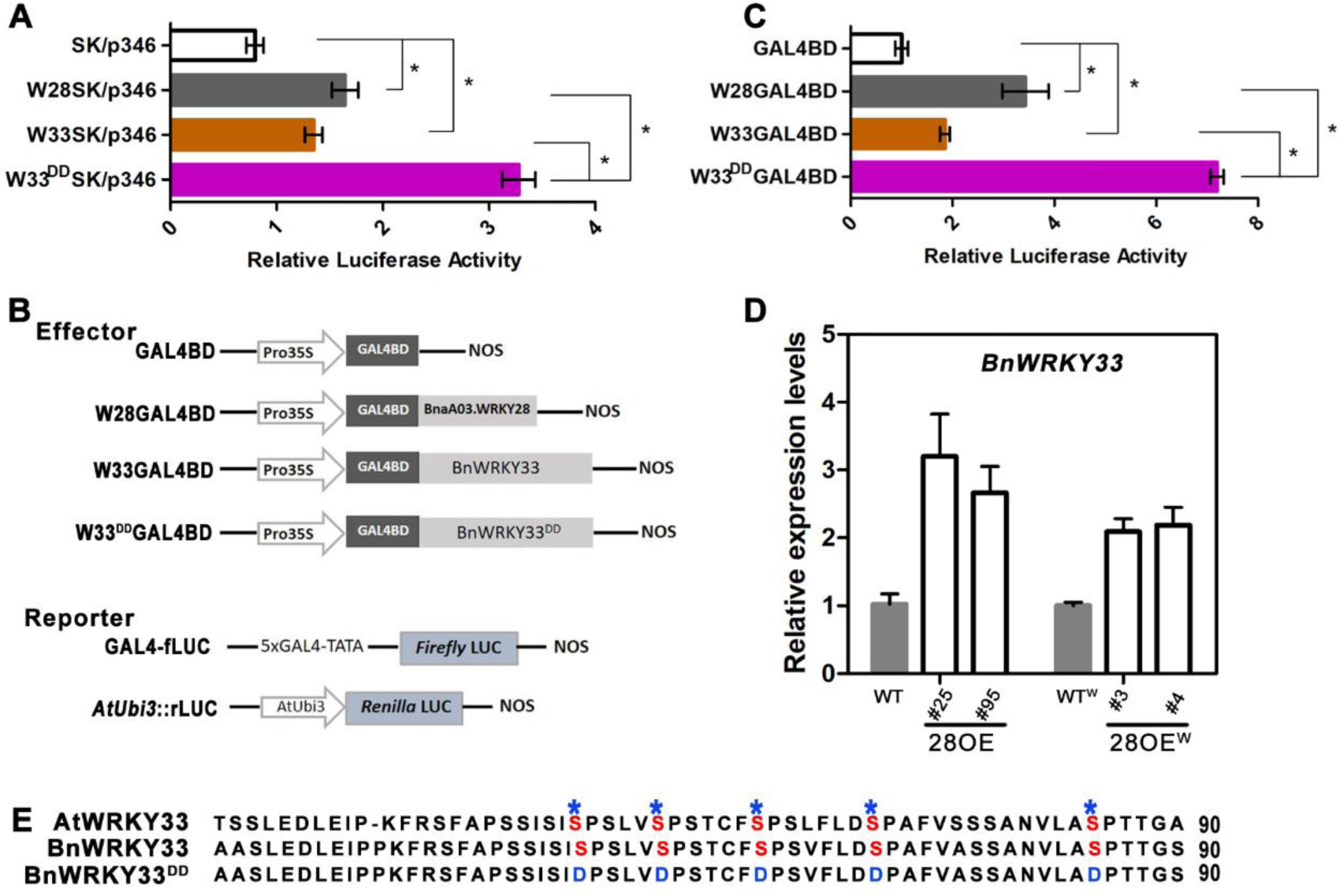
The N-terminal Modification of BnWRKY33 is Necessary to Increase the Transcriptional Activity of BnWRKY33. **(A)** Dual luciferase assays using the pGreenII vector set showing the relative luciferase activity of BnaA03.WRKY28, BnWRKY33, and BnWRKY33^DD^ (DD indicates that five Ser residues in the N-terminus of BnWRKY33 were all mutated to Asp) when binding to the *BnWRKY33* promoter. Data are shown as means ± SD (*n* = 3). Asterisks indicate significant differences (t-test, \**P* < 0.05). **(B)** Schematic diagram of another set of vectors to detect the transcriptional activity of BnaA03.WRKY28, BnWRKY33, and BnWRKY33^DD^. The transcription factor fused with the yeast GAL4 binding domain (GAL4BD) was used as an effector; GAL4-fLUC was used as a reporter, and *AtUbi:rLUC* was used as an internal control. **(C)** Dual luciferase assays showing comparisons of the transcriptional activities of these transcription factors. Data are shown as means ± SD (*n* = 3). Asterisks indicate significant differences (t-test, \**P* < 0.05). **(D)** qRT-PCR was performed to detect the transcript abundance of *BnWRKY33* in *BnaA03.WRKY28* overexpression (28OE) lines and wild-type plants without inoculation with fungi. Superscript W in WT^W^ and 28OE^W^ indicate wild-type rapeseed varieties Westar and *BnaA03.WRKY28* overexpression in Westar background, respectively. Data are shown as means ± SD (*n* = 3). **(E)** A diagram of the N-terminus of BnWRKY33 showing the positions of five conserved Ser residues according to AtWRKY33. BnWRKY33^DD^, the five Ser residues were all mutated to Asp.

In Arabidopsis, it is reported that WRKY33 can be phosphorylated by MPK3 and MPK6. The phosphorylation modification does not alter the binding activity of WRKY33 to downstream genes but can activate the activating activity of WRKY33 (Mao et al., 2011). In our study, both the measure of LUC activity and the expression of *BnWRKY33* in *BnaA03.WRKY28* overexpression lines were conducted in the static state (without inoculation); that was, BnWRKY33 may be in the passivation (unphosphorylated) state. Therefore, we speculated that BnWRKY33, as a positive disease resistance factor, could be activated by phosphorylation after infection with Sclerotinia in *Brassica napu*s. A cluster of five Ser residues in the N-terminus of AtWRKY33 is MPK3/6 phosphorylation site (Mao et al., 2011), the domain of which is conserved in BnWRKY33 (Figure 3E). By point-directed mutagenesis, these five Ser residues were all mutated to Asp (*BnWRKY33^DD^*). *BnWRKY33^DD^* was cloned into the pGreenII 62-SK and pGAL4BD vectors. As expected, in dual-luciferase transient transcriptional activity assays, the relative LUC activity of cotransformed *Pro35S:BnWRKY33^DD^* with *p346:LUC* (W33^DD^SK/p346) was much higher than that of W33SK/p346 or W28SK/p346 (Figure 3A). Similar results were found in the comparison of W33^DD^GAL4BD and W33GAL4BD or W28GAL4BD (Figure 3B), suggesting that the activation capacity of phosphorylation-activated BnWRKY33 was stronger than that of BnaA03.WRKY28. These results indicate that BnWRKY33 is activated by phosphorylation to enhance its transcriptional activity.

### MAPK Signal Cascade: BnMKK4 Interacts with BnMPK3 and Phosphorylates BnMPK3, and BnMPK3 Phosphorylates BnWRKY33 and Promotes its Transactivation Activity by Interacting with BnWRKY33

There were two homologous copies of *AtMPK3* in rapeseed, located on chromosomes A06 and C03, both of which could be significantly induced by *S. sclerotiorum* (Supplemental Figure 5A and 5B). In contrast, *AtMKK4* and *AtMKK5* are functionally redundant (Meng and Zhang, 2013), and both have multiple homologs in rapeseed (data not shown). Nevertheless, *BnaA03.MKK4*, one of the homologs of *AtMKK4*, was reported to be involved in low-temperature and salt stress response (Zhang et al., 2012), and could be induced by *S. sclerotiorum* (Supplemental Figure 5C). In the yeast two-hybrid (Y2H) assays, we found that BnaA03.MKK4 and BnWRKY33 interacted with BnaA06.MPK3 or BnaC03.MPK3 (Figure 4A and 4B). Next, we performed pull-down assays to confirm the interactions between these proteins *in vitro*. His-tagged BnaA03.MKK4 and BnWRKY33 were pulled down by GST-BnaA06.MPK3 or GST-BnaC03.MPK3, but not by GST alone (Supplemental Figure 6A and 6B). The interactions were further confirmed using coimmunoprecipitation (CoIP) assays in which BnaA03.MKK4–FLAG, BnWRKY33–FLAG BnaA06.MPK3-GFP, and BnaC03.MPK3-GFP fusion proteins were coexpressed in *Nicotiana benthamiana* leaves. BnaA03.MKK4–FLAG and BnWRKY33–FLAG coimmunoprecipitated with BnaA06.MPK3-GFP or BnaC03.MPK3-GFP instead of the negative control (Figure 4C and 4D). Taken together, both *in vitro* and *in vivo* data indicate that BnaA03.MKK4 and BnWRKY33 physically interact with BnaA06.MPK3 or BnaC03.MPK3.

**Figure 4.**
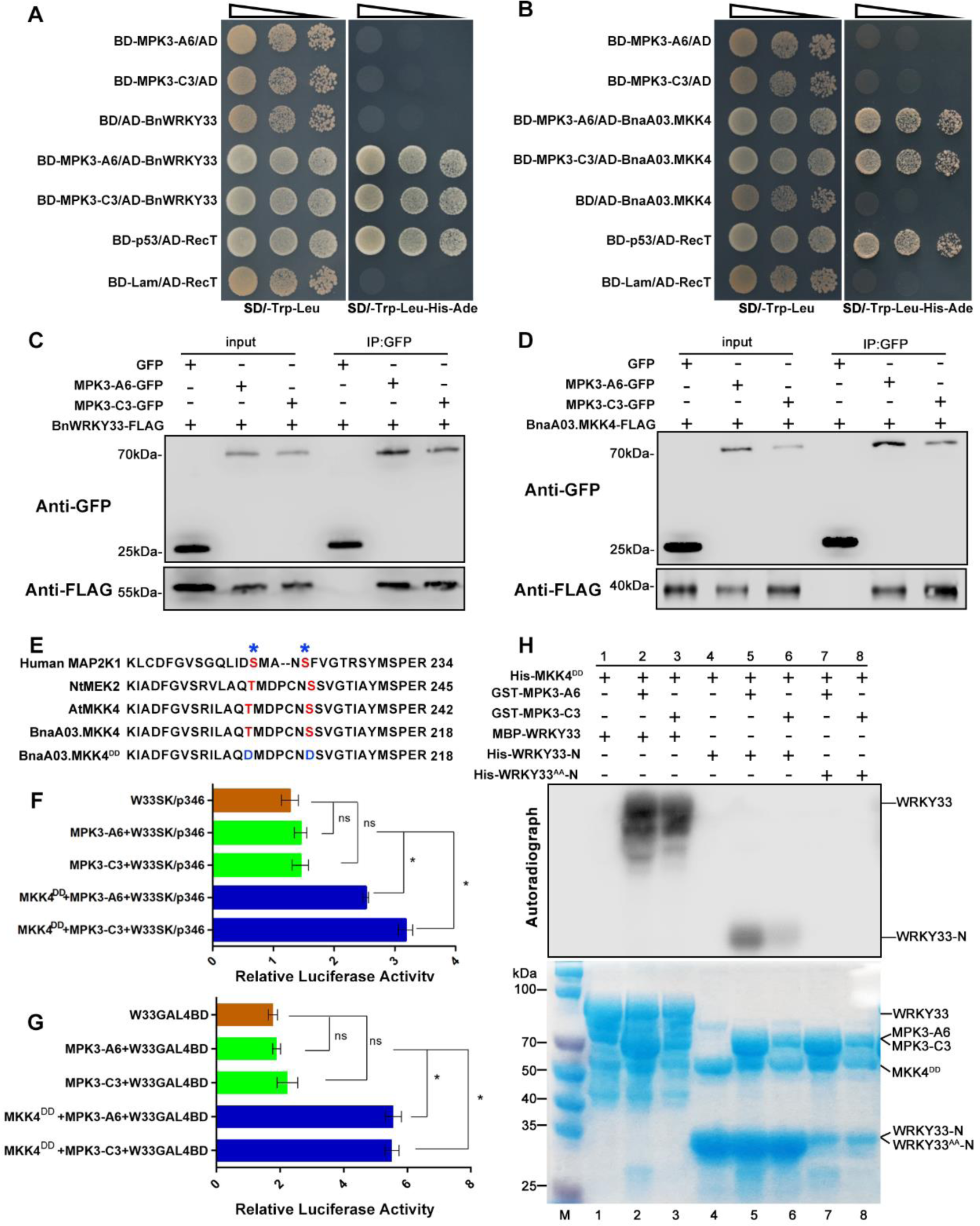
BnMKK4-BnMPK3 Module Phosphorylates BnWRKY33 and Promotes its Transactivation Activity Burst by Interacting with BnWRKY33. **(A)** and **(B)** Yeast two-hybrid assays indicate that BnWRKY33 interacts with BnaA06.MPK3/BnaC03.MPK3, and BnaA03.MKK4 interacts with BnaA06.MPK3/BnaC03.MPK3. The yeast cells were cultured on SD/-Trp-Leu (selective dropout medium without tryptophan and leucine) or SD/-Trp-Leu-His-Ade (selective dropout medium without tryptophan, leucine, histidine, and adenine). AD, pGADT7/activation domain; BD, pGBKT7/binding domain. BD- p53/AD-RecT, positive control; BD-Lam/AD-RecT, negative control. The oblique triangles above the images indicate 10^0^, 10^-1^, and 10^-2^ concentration gradient of Yeast cells. **(C)** and **(D)** Coimmunoprecipitation (CoIP) assays indicate that BnWRKY33 interacts with BnaA06.MPK3/BnaC03.MPK3, and BnaA03.MKK4 interacts with BnaA06.MPK3/BnaC03.MPK3 in planta. Fusion proteins were coexpressed in tobacco leaves. Total proteins were extracted (Input) and immunoprecipitated (IP) with GFP beads. The immunoblot assays were performed with anti-GFP and anti-FLAG antibodies. **(E)** A diagram of the conserved Ser/Thr in the activation loop of BnaA03.MKK4 by alignment to the homologous genes in other species. BnaA03.MKK4^DD^, the conserved Ser/Thr were all mutated to Asp. **(F)** and **(G)** Dual luciferase assays show that the activated MAPK cascade can significantly enhance the transcriptional activity of BnWRKY33. BnWRKY33 (W33), BnaA06.MPK3 (MPK3-A6), BnaC03.MPK3 (MPK3-C3), and BnaA03.MKK4^DD^ (MKK4^DD^) were introduced into pGreenII 62-SK (SK) and pGAL4BD as effectors, respectively. Data are shown as means ± SD (*n* = 3). Asterisks indicate significant differences (t-test, ns, *P* > 0.05, \**P* < 0.05). **(H)** BnWRKY33 phosphorylation mediated by activated BnaA06.MPK3 (MPK3- A6) and BnaC03.MPK3 (MPK3-C3) *in vitro*. MPK3-A6 and MPK3-C3 were activated with constitutively active BnaA03.MKK4^DD^ (MKK4^DD^). WRKY33-N, the N-terminus of BnWRKY33 containing the five conserved Ser residues; WRKY33^AA^-N, the five conserved Ser residues in the N-terminus of BnWRKY33 were all mutated to Ala. The phosphorylation reactions were incubated in protein kinase buffer containing [γ-^32^P] ATP. The phosphorylation signal was detected by autoradiography (top panels) and the protein inputs were assessed by Coomassie Brilliant Blue (CBB) staining (bottom panels).

Therefore, we performed phosphorylation assays to determine whether BnWRKY33 can be phosphorylated by MAPK cascade *in vitro*. According to previous research, constitutively active BnMKK4 was used in phosphorylation test assays (Ren et al., 2002; Mao et al., 2011). We generated an active mutant of BnaA03.MKK4 named BnaA03.MKK4^DD^, by mutating the conserved Ser/Thr in the activation loop ((S/T)XXXXX(S/T)) of plant MAPKKs to Asp (Yang et al., 2001) (Figure 4E). In the autoradiogram, recombinant GST-BnaA06.MPK3 and GST-BnaC03.MPK3 strongly phosphorylated BnWRKY33 after activation by constitutively active BnaA03.MKK4^DD^ (Figure 4H). As mentioned above, a cluster of five Ser residues in the N-terminus of BnWRKY33 may be the phosphorylation sites. The phosphorylation signal could also be detected when the truncated N-terminal containing these five Ser was used as a substrate, but if the five Ser residues were mutated into Ala (named WRKY33^AA^-N), it cannot be phosphorylated by BnMPK3 (Figure 4H). These results demonstrate that BnWRKY33 is a substrate of the BnaA06.MPK3/BnaC03.MPK3-mediated cascade.

In dual-luciferase transient transcriptional activity assays, we found that without activation by the constitutively active BnaA03.MKK4^DD^, neither BnaA06.MPK3 nor BnaC03.MPK3, was able to enhance the activity of BnWRKY33. Nevertheless, the relative LUC activity significantly increased when the MAPK cascade was activated (Figure 4F and 4G). Overall, these results suggest that BnaA03.MKK4 interacts with, phosphorylates and activates BnaA06.MPK3/BnaC03.MPK3, while BnaA06.MPK3/BnaC03.MPK3 interacts with, phosphorylates and activates BnWRKY33.

### BnaA03.MKK4 and BnaA06.MPK3/BnaC03.MPK3 Contribute to S. sclerotiorum Resistance in Brassica napus

To determine whether *BnaA03.MKK4* and *BnaA06.MPK3/BnaC03.MPK3* function in Sclerotinia resistance, we generated transgenic rapeseed with *BnaA06.MPK3/BnaC03.MPK3* overexpression driven by CaMV35S, *BnMPK3* knockout mediated by CRISPR/Cas9, and *BnaA03.MKK4* overexpression driven by CaMV35S.

Three independent *BnaA06.MPK3* overexpression lines, as well as three independent *BnaC03.MPK3* overexpression lines with more than 10-fold increase in expression (Figure 5A and 5B), and homozygous mutants with three editing types in the genome region of *BnaA06.MPK3* and *BnaC03.MPK3* mediated by CRISPR/Cas9 (Figure 5C) were established for comparison with wild-type Jia9709. The conserved amino acid motif TDY/TEY of MAPKs is phosphorylated by MKKs (Ichimura et al., 2002; Meng and Zhang, 2013), and the common docking (CD) site located in the carboxyl-terminal of MAPKs includes two adjacent acidic residues (D and E) that are crucial for the interaction with MKKs (Ichimura et al., 2002; Doczi et al., 2012) (Figure 5C). After 48 h of inoculation with *S. sclerotiorum*, it was determined that the lesion areas of both *BnaA06.MPK3* and *BnaC03.MPK3* overexpression lines were smaller than those of WT. However, the knockout lines showed a decrease in pathogen resistance (Figure 5D and 5E). The expression of *BnPAD3* and *BnCYP71A13* in *BnMPK3* overexpression lines were much higher than those in WT, and remained at a high level with the invasion of pathogens. In BnMPK3 knockout lines, the expression of these genes did not change much and were at a very low level (Figure 5F and 5G). In addition, we examined the phenotype of *BnaA03.MKK4* overexpression lines (MKK4-OE) with different expression levels after inoculation (Supplemental Figure 7A and 6B). It was found that MKK4-OE lines were more resistant against Sclerotinia than WT (Supplemental Figure 7C). In contrast to wild-type plants, the MKK4-OE lines showed increased transcript accumulation of *BnPAD3* and *BnCYP71A13* (Supplemental Figure 7D and 6E). These results show that in *Brassica napus*, BnaA03.MKK4 and BnaA06.MPK3/BnaC03.MPK3 are activated to resist the invasion of pathogen when plants suffered from *S. sclerotiorum*.

**Figure 5.**
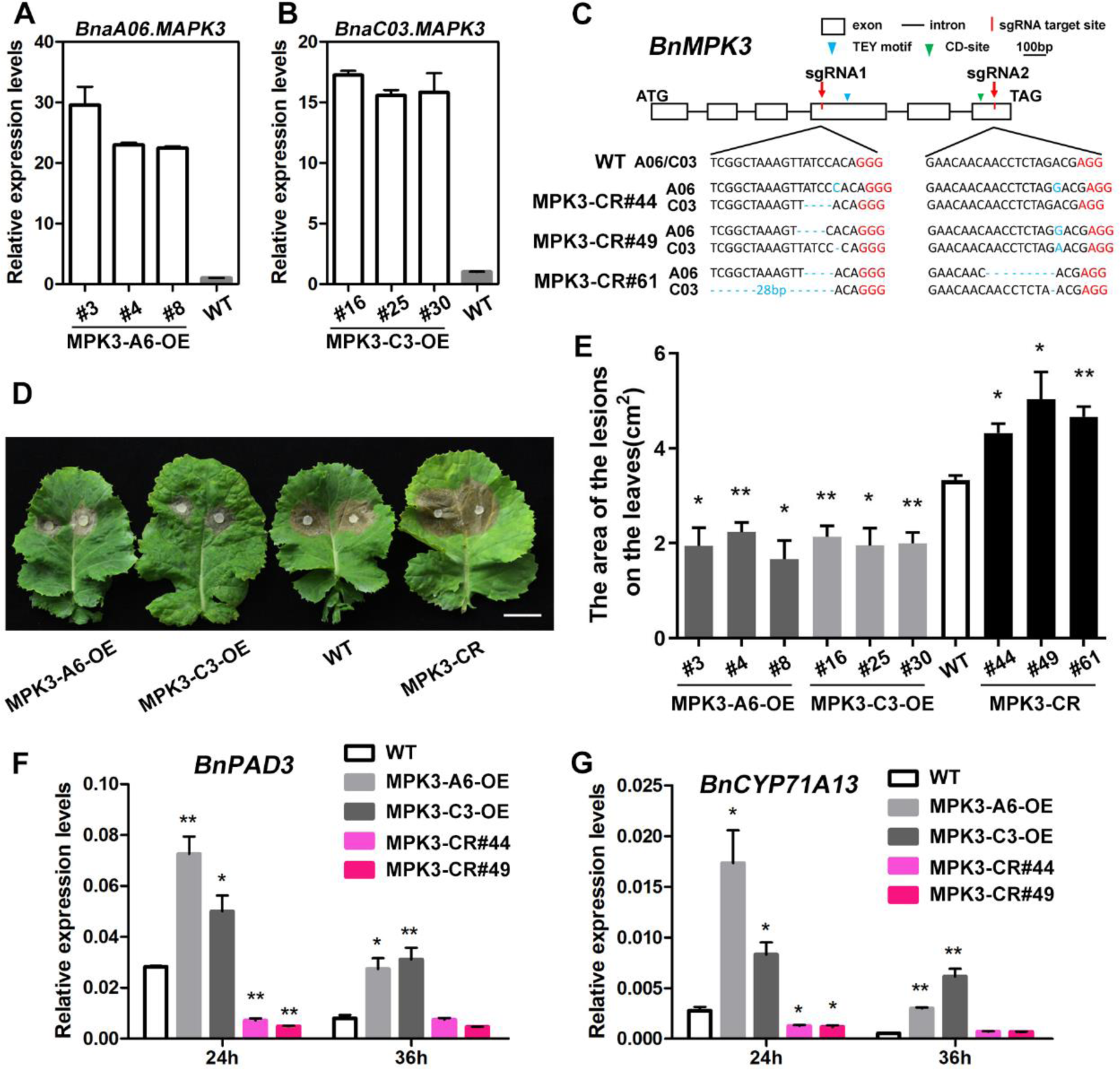
*BnMPK3* Positively Contributes to *S. sclerotiorum* Resistance in *Brassica napus.* **(A)** and **(B)** *BnaA06.MPK3* (MPK3-A6) overexpression (OE) lines and *BnaC03.MPK3* (MPK3-C3) OE lines establishment by qRT-PCR analysis. The expression in WT as control. Data are shown as means ± SD (*n* = 3). **(C)** A schematic diagram to show the characterization of the *BnMPK3* exon/intron sequence and the target sites of sgRNAs. The edit types of three independent homozygous mutants (MPK3-CR#44, 49, and 61) are shown. The protospacer-adjacent motifs (PAM) are marked in red. TEY motif, phosphorylated by MKKs; CD-site, interacts with MKKs; ATG, start codon; TAG, stop codon. **(D)** Photographs of the lesions of detached leaves from *BnMPK3* transgenic lines and WT plants (Jia 9709) in approximately 4 weeks when inoculated with 7 mm *S. sclerotiorum* hyphae agar block at 22°C for 48 h. Scale bar, 2 cm. **(E)** Statistical analysis of the lesion areas of *BnMPK3* transgenic lines and WT plants after 48 h inoculation. Data are shown as means ± SD (*n* = 3). Asterisks indicate significant differences compared with WT (t-test, \**P* < 0.05, \*\**P* < 0.01). **(F)** and **(G)** Relative expression levels of two camalexin biosynthesis-related genes, *BnPAD3* and *BnCYP71A13*, in *BnMPK3* transgenic lines and WT plants after *S. sclerotiorum* inoculation for 24 h or 36 h. Data are shown as means ± SD (*n* = 3). Asterisks indicate significant differences compared with WT at corresponding time points (t-test, \**P* < 0.05, \*\**P* < 0.01).

### As a Co-factor, BnaA09.VQ12 Interacts with BnaA03.WRKY28 to Form a Protein Complex to Negatively Regulate Sclerotinia Resistance

It is reported that VQ proteins usually regulate gene expression by interacting with WRKY TFs (Cheng et al., 2012). AtVQ12 plays a negative role in the resistance of necrotrophic fungal pathogen *B. cinerea* (Wang et al., 2015). There were two homologous copies of *BnVQ12* (*BnaA09.VQ12* and *BnaC08.VQ12*) in rapeseed, between which *BnaA09.VQ12* could be markedly induced by *S. sclerotiorum*, and the induction trend was similar to that of *BnaA03.WRKY28* (Figure 6A). We first tested the interaction between BnaA03.WRKY28 and BnaA09.VQ12 in Y2H assays. We fused BnaA03.WRKY28 with the DNA binding domain (BD) of GAL4 (BD-WRKY28), and found that the full-length BnaA03.WRKY28 itself had transcription-activating activity to activate reporter genes in yeast (Figure 6B). Then different forms of truncated domains of BnaA03.WRKY28 were fused into BD vector, and it was found that the activation domain was located in the first 86 amino acids (aa) at the N-terminal (Figure 6B). Full-length *BnaA09.VQ12* and *BnA01.VQ22* were cloned into the prey vector (AD). It was discovered that BnaA03.WRKY28 with a deleted activation domain (WRKY28 Δ1-86) interacted with BnaA09.VQ12 but not BnA01.VQ22 (Figure 6C), indicating the specificity of the BnaA03.WRKY28-BnaA09.VQ12 interaction. Furthermore, Y2H assays suggested that BnaA09.VQ12 interacted with BnaA03.WRKY28 through the 164-227 aa domain of BnaA03.WRKY28, which is the DNA binding domain containing the WRKY motif and C_2_H_2_ zinc finger (Figure 6D). The biomolecular florescence complementation (BiFC) assays showed that BnaA09.VQ12 interacted with BnaA03.WRKY28 in the nucleus (Figure 6E). However, subcellular localization analysis by transient expression of *Pro35S:BnaA03.WRKY28:GFP* and *Pro35S:BnaA09.VQ12:GFP* in Arabidopsis protoplasts showed that BnaA03.WRKY28 was localized in the nucleus, while BnaA09.VQ12 could be found in the cytoplasm and nucleus (Supplemental Figure 8). In addition, GST pull-down assays were conducted to confirm the interactions *in vitro*. GST-BnaA03.WRKY28 and GST-BnaA03.WRKY28 164-227 (the 164-227 aa domain of BnaA03.WRKY28), but not the GST control, pulled down His-BnaA09.VQ12 (Figure 6F). CoIP assays further showed that BnaA09.VQ12-FlAG could interact with BnaA03.WRKY28-GFP and even with BnaA03.WRKY28 164-227 (Figure 6G). Thus, the co-factor BnaA09.VQ12 forms a complex with BnaA03.WRKY28 in the nucleus by binding to the DNA binding domain of the transcription factor BnaA03.WRKY28.

**Figure 6.**
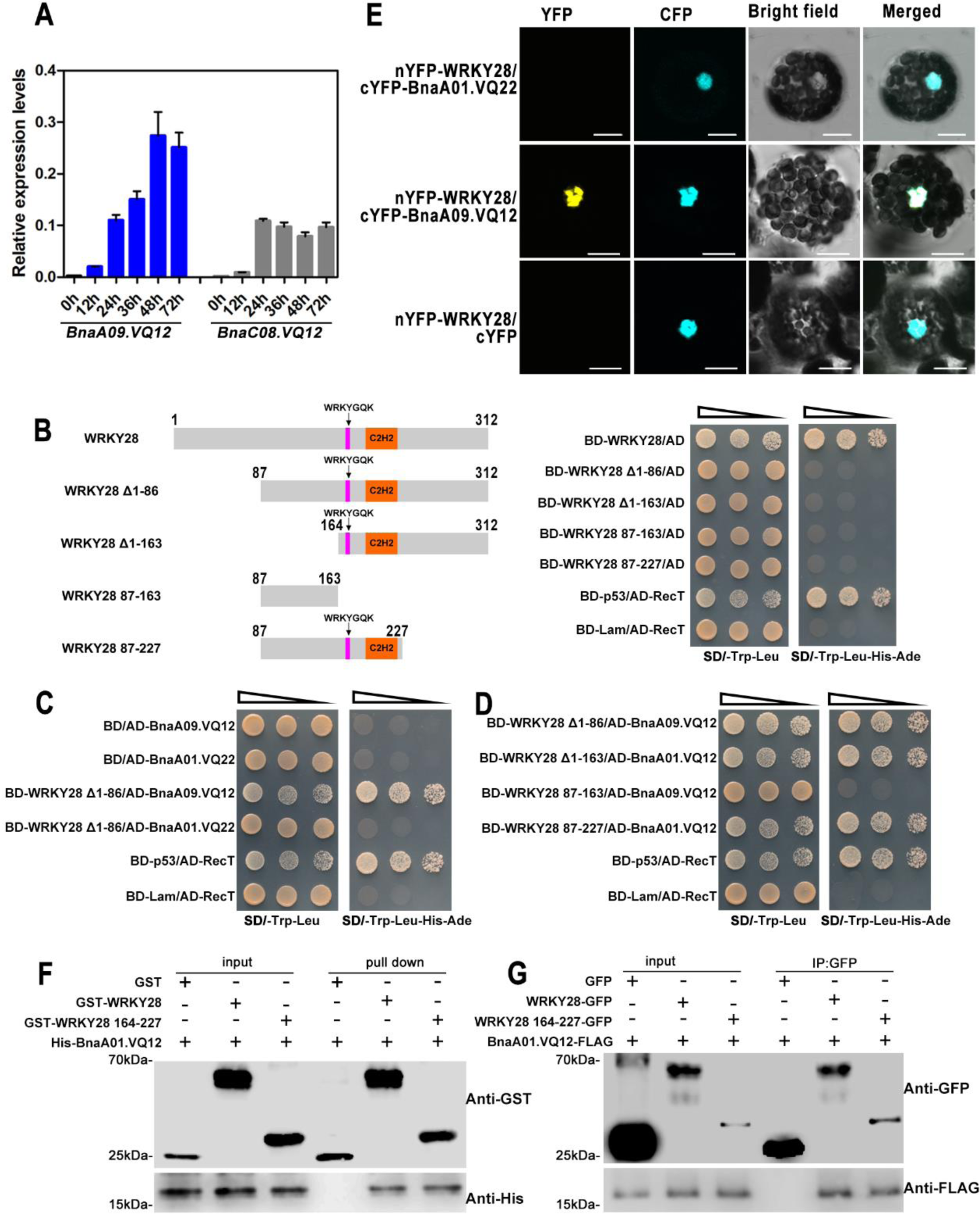
BnaA03.WRKY28 Interacts with BnaA09.VQ12 *in vitro* and *in vivo.* **(A)** Induced expression profiles of two copies (A09 and C08) of *BnVQ12* were identified at 0 h, 12 h, 24 h, 36 h, 48 h, and 72 h after inoculation with *S. sclerotiorum*. The latest fully unfolded leaves of rapeseed (Jia 9709) were used for infection in approximately 4 weeks. Data are shown as means ± SD (*n* = 3). **(B)** Diagrams of the full-length and truncated BnaA03.WRKY28 constructs with specific deletions (left), and identification of the self-activation region of BnaA03.WRKY28 in yeast cells (right). Full-length of BnaA03.WRKY28 contains 312 amino acids (aa). The magenta rectangles denote WRKYGQK motif; the orange rectangles denote the zinc-finger motif. BD-p53/AD-RecT, positive control; BD-Lam/AD-RecT, negative control. **(C)** Identification of the interaction between BnaA03.WRKY28 and BnaA09.VQ12 in yeast cells. BnaA03.WRKY28 interacts with BnaA09.VQ12, but not with BnaA01.VQ22. **(D)** Identification of the specific domain of BnaA03.WRKY28 that interacts with BnaA09.VQ12. **(E)** Bimolecular fluorescence complementation (BiFC) assays to confirm the BnaA03.WRKY28 and BnaA09.VQ12 interaction in *Arabidopsis* protoplasts. BnaA01.VQ22 and cYFP served as negative controls. Scale bars, 10 μm. **(F)** *In vitro* pull-down assays showing the direct interactions of BnaA03.WRKY28, especially the DNA binding domain of BnaA03.WRKY28 (164-227 aa) with BnaA09.VQ12. GST, GST-WRKY28, or GST-WRKY28 164- 227 was immobilized on GST beads and incubated with His-BnaA09.VQ12 protein. The immunoblot assays were performed using anti-GST and anti-His antibodies. **(G)** CoIP assays with BnaA03.WRKY28-GFP and BnaA09.VQ12-FLAG coexpressed in *N. benthamiana* leaves. Proteins were extracted (Input) and immunoprecipitated (IP) with GFP beads. The immunoblot assays were performed using anti-GFP and anti-FLAG antibodies.

To characterize the biological function of *BnaA09.VQ12* in response to pathogen infection, three independent *BnaA09.VQ12* overexpression lines and three independent loss-of-function homozygous mutants with two copies of *BnVQ12* edited by the CRISPR/Cas9 system were obtained (Figure 7A and 7B). At 48 h post-inoculation, the necrotic symptoms of *BnaA09.VQ12* overexpression lines were more severe than those of WT, whereas *BnVQ12* knockout lines had fewer symptoms compared with WT (Figure 7C and 7D). After *S. sclerotiorum* infection, the transcripts of *BnPAD3* and *BnCYP71A13* in *BnaA09.VQ12* overexpression lines were significantly reduced compared with WT, but increased in *BnVQ12* knockout lines. In detail, in WT plants, the camalexin synthesis-related genes were significantly induced at 24 h after incubation and decreased significantly at 36 h. However, in *BnaA09.VQ12* overexpression lines, the induced expression of these genes was no longer obvious at 24 h, and was still at a low-expression level at 36 h. In *BnVQ12* knockout lines, the induced expression levels of these genes were similar to those of WT at 24 h, and the expression levels of these genes decreased at 36 h, but were significantly higher than that of WT (Figure 7E and 7F). These results demonstrate that *BnaA09.VQ12*, similar to its partner *BnaA03.WRKY28*, negatively contributed to *S. sclerotiorum* resistance.

**Figure 7.**
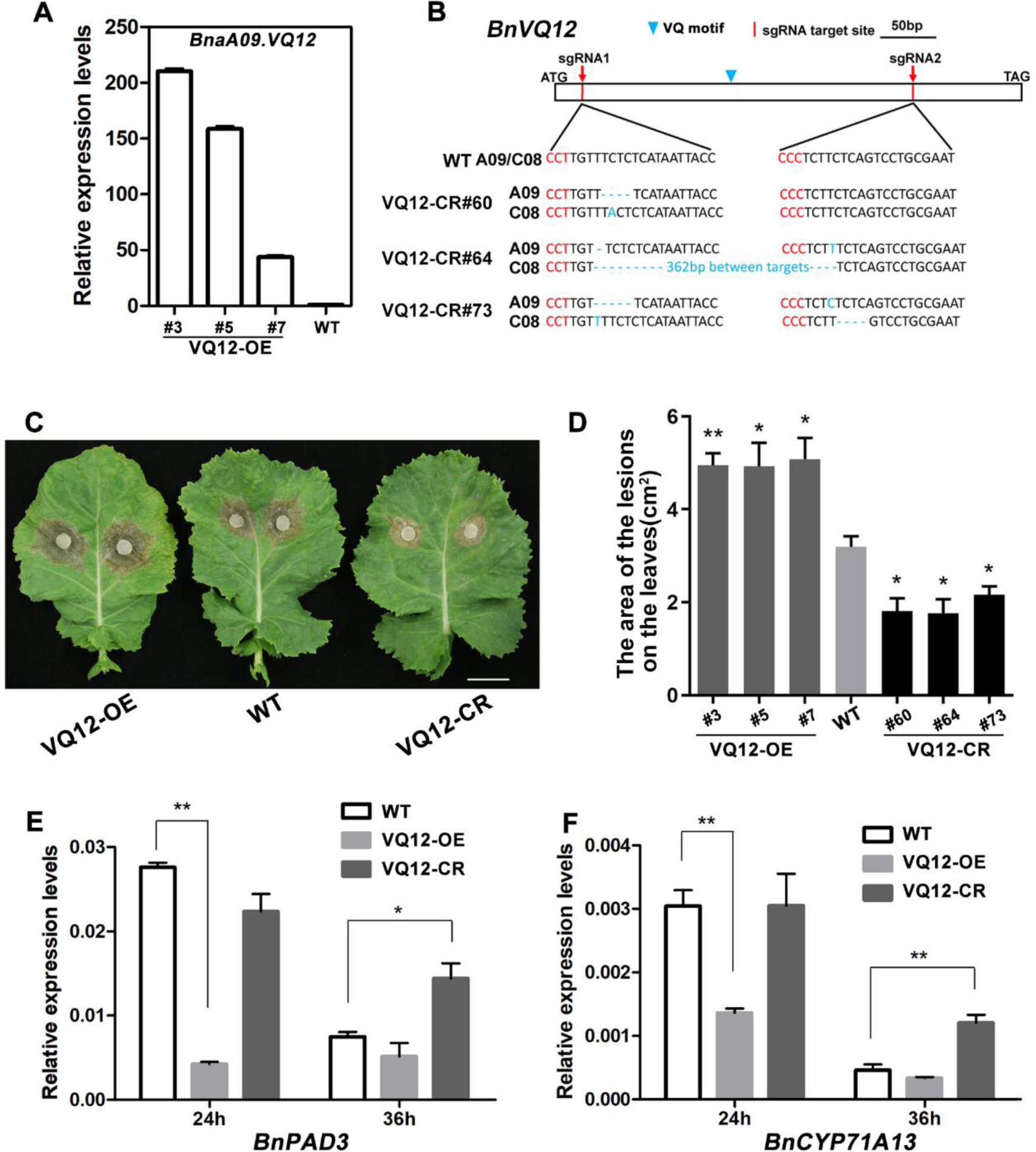
*BnaA09.VQ12* Plays a Negative Role in *S. sclerotiorum* Resistance in *Brassica napus.* **(A)** *BnaA09.VQ12* overexpression (VQ12-OE) lines establishment by qRT-PCR analysis. The expression of *BnaA09.VQ12* in WT plants is the control. Data are shown as means ± SD (*n* = 3). **(B)** A schematic diagram to show the characterization of the *BnVQ12* sequence and the target sites of sgRNAs. The edit types of three independent homozygous mutants (VQ12-CR#60, 64, and 73) are shown. The protospacer-adjacent motifs (PAM) are marked in red. VQ, conserved motif of VQ proteins. **(C)** and **(D)** *BnVQ12* transgenic homozygous lines and WT (Jia 9709) were imaged after 48 h inoculation with *S. sclerotiorum* and the lesion areas were statistically analyzed. Scale bar, 2 cm. Data are shown as means ± SD (*n* = 3). Asterisks indicate significant differences compared with WT (t-test, \**P* < 0.05, \*\**P* < 0.01). **(E)** and **(F)** Relative expression levels of *BnPAD3*/*BnCYP71A13* in *BnVQ12* transgenic lines and WT plants after *S. sclerotiorum* inoculation for 24 h or 36 h. Data are shown as means ± SD (*n* = 3). Asterisks indicate significant differences compared with WT (t-test, \**P* < 0.05, \*\**P* < 0.01).

### The Formation of WRKY28-VQ12 Complex Promotes BnaA03.WRKY28 to Induce a Higher Affinity for the *BnWRKY33* Promoter than BnWRKY33

We tracked the degree increase of lesions on rapeseed leaves when infected by *S. sclerotiorum* within 0-84 hours, and roughly plotted the increase rate of lesions every 3 h (Figure 8A). It was determined that the increase rate of lesions was at the slowest level within 15-24 h after inoculation, which was the period when *BnWRKY33* was strongly induced (Figure 8B). However, from 36 h after inoculation, the degree of lesions continued to rapidly increase, and may reach a peak within 42-48 h, which was also the stage in which *BnaA03.WRKY28* and *BnaA09.VQ12* were dramatically induced (Figure 1A, Figure 6A), whereas the expression of *BnWRKY33* decreased sharply.

**Figure 8.**
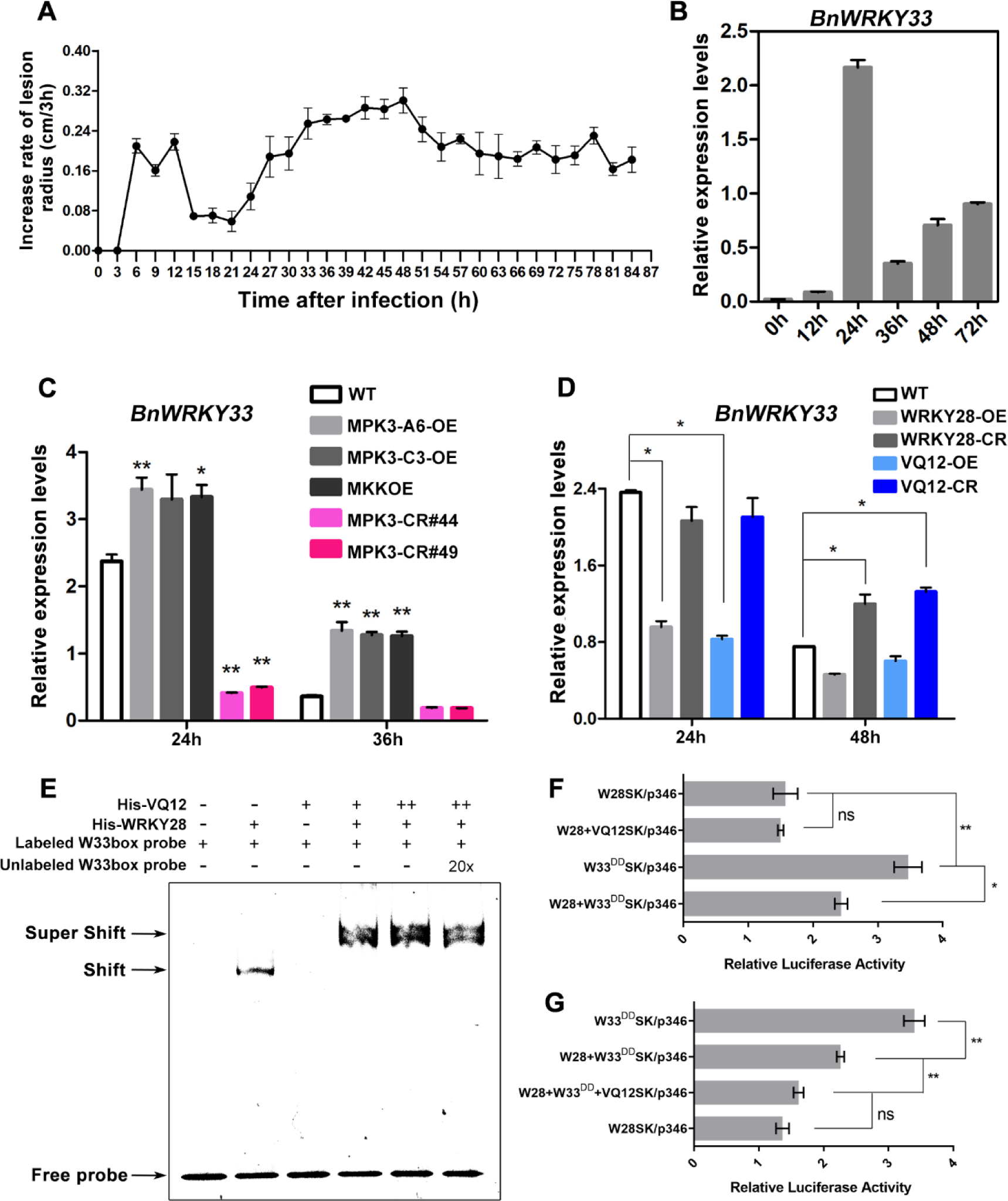
The BnaA09.VQ12-BnaA03.WRKY28 Protein Complex Inhibits the Transcriptional Burst of *BnWRKY33* by Competitive Binding to *BnWRKY33* Promoter. **(A)** A diagram of the increase rate of lesion radius when rapeseed leaves were infected by *S. sclerotiorum* within 0-84 h. The lesions were measured every 3 h. Data are shown as means ± SD (*n* = 3). **(B)** Induced expression profile of *BnWRKY33* was identified at 0 h, 12 h, 24 h, 36 h, 48 h, and 72 h after inoculation with *S. sclerotiorum* in WT. Data are shown as means ± SD (*n* = 3). **(C)** and **(D)** Relative expression levels of *BnWRKY33* in *BnMPK3/ BnaA03.MKK4* transgenic lines and WT plants **(C)**, in *BnaA03.WRKY28/ BnaA09.VQ12* transgenic lines and WT plants **(D)** after *S. sclerotiorum* inoculation for 24 h and 36/48 h. Data are shown as means ± SD (*n* = 3). Asterisks indicate significant differences compared with WT at corresponding time points (t-test, \**P* < 0.05, \*\**P* < 0.01). **(D)** EMSA assays showing the effect of BnaA09.VQ12-BnaA03.WRKY28 protein complex on the binding capacity of BnaA03.WRKY28 to the *BnWRKY33* promoter. **(F)** and **(G)** Dual luciferase assays show that BnaA03.WRKY28 and BnWRKY33 compete to bind to the promoter of *BnWRKY33* **(F)**, and BnaA09.VQ12 helps BnaA03.WRKY28 preferentially bind to the promoter of *BnWRKY33* **(G)**. Data are shown as means ± SD (n = 3). Asterisks indicate significant differences (t-test, \**P* < 0.05, \*\**P* < 0.01). At least three independent experiments were performed with similar results.

After infection, the expression levels of *BnWRKY33* in *BnMPK3*/*BnaA03.MKK4* overexpression lines were significantly higher than that in WT (at 24 h and 36 h), but dramatically decreased in *BnMPK3* knockout lines, especially at 24 h (Figure 8C). In *BnMPK3* knockout lines, the expression of *BnWRKY33* changed little and was at a very low level. However, the expression of *BnWRKY33* in *BnaA03.WRKY28*/*BnaA09.VQ12* overexpression lines was significantly decreased compared with that in WT, especially at 24 h after inoculation, when the expression of *BnWRKY33* reached its peak in WT (Figure 8D). However, 24 h after inoculation, the expression of *BnWRKY33* in *BnaA03.WRKY28* knockout and *BnaA09.VQ12* knockout plants were similar to that of WT, but at 48 h, the expression of *BnWRKY33* in *BnaA03.WRKY28*/*BnaA09.VQ12* knockout plants was significantly higher than that of WT. These results suggest that BnaA03.MKK4-BnaA06.MPK3/BnaC03.MPK3 module promotes a transcriptional burst of *BnWRKY33*, whereas the BnaA03.WRKY28-BnaA09.VQ12 complex may inhibit the expression of *BnWRKY33*. Because both BnaA03.WRKY28 and BnWRKY33 could bind to the promoter of *BnWRKY33*, and the transcriptional activity of BnaA03.WRKY28 was lower than that of BnWRKY33, we speculated that BnaA03.WRKY28 may inhibit the binding of BnWRKY33 to the *BnWRKY33* promoter.

Firstly, EMSA assays were conducted to examine the effects of the interaction between BnaA09.VQ12 and BnaA03.WRKY28 on the binding of BnaA03.WRKY28 to the promoter of *BnWRKY33* (Figure 8E). In the absence of BnaA09.VQ12, BnaA03.WRKY28 bound to *BnWRKY33* and formed a specific shifted band, whereas in the absence of BnaA03.WRKY28, BnaA09.VQ12 failed to target *BnWRKY33.* When BnaA09.VQ12 was added to the mixture of BnaA03.WRKY28 and DNA probe, a specific super-shifted band was produced, whose intensity was enhanced when more BnaA09.VQ12 was added. These results suggest that the formation of BnaA03.WRKY28-BnaA09.VQ12 complex stimulates the binding activity of BnaA03.WRKY28 to *BnWRKY33*.

Next, in dual-luciferase transient transcriptional activity assays, we found that the relative LUC activity displayed an obvious trend when W28SK, W33^DD^SK, and VQ12SK were used as different effector combinations. The relative LUC activity of W28SK as an effector alone was similar to that of W28SK and VQ12SK as effectors simultaneously (Figure 8F), indicating that the binding of BnaA09.VQ12 and BnaA03.WRKY28 did not change the transcriptional activity of BnaA03.WRKY28. When W28SK and W33^DD^SK were contained at the same time, the relative LUC activity was between the relative LUC activities when they were used effector alone (Figure 8F). However, when VQ12SK was reintroduced into the mixture of W28SK and W33^DD^SK, the relative LUC activity tended to be similar to that of W28SK as an effector alone (Figure 8G). Based on these data, we believe that BnaA03.WRKY28 and BnWRKY33 compete to bind to the promoter of *BnWRKY33* when they exist simultaneously, but the present of BnaA09.VQ12 makes BnaA03.WRKY28 have stronger binding capacity the *BnWRKY33* promoter than BnWRKY33.

### BnaA03.WRKY28 Contributes to the Promotion of Axillary Bud Activity and Shoot Branching

Two or more branches were produced in one leaf axil and more higher-order branches were formed in the *BnaA03.WRKY28* overexpression lines (Figure 9A, Supplemental Figure 9A to 9C). When the number of branches and buds developing in the leaf axils was quantified, it was found that there was at most one branch formed in one leaf axil and secondary branching was the highest-order branching in WT. In the *BnaA03.WRKY28* overexpression lines, two or more branches formed in one leaf axil and quaternary branches were very common, and even quindary branches could be found (Figure 9B). We also observed that at the late stage of rapeseed growth, when the siliques of WT matured, the siliques were light yellow without cauline leaves, whereas the overexpression lines showed a certain degree of green late ripening, with green siliques and many cauline leaves (Figure 9A, Supplemental Figure 9C). By GUS staining analysis, we found that *BnaA03.WRKY28* was expressed in the latest true leaves and the axils at the seedling stage, and also expressed in the leaf axil at the flowering stage (Figure 9C). Moreover, among the DEGs of RNA-seq, the expression levels of genes involved in meristem growth and auxin polar transport were significantly increased in the *BnaA03.WRKY28* overexpression lines compared to those in WT, whereas the expression levels of genes involved in auxin influx, cellular response to auxin stimulus, and auxin biosynthetic process were significantly reduced (Supplemental Figure 9D and 9E, Supplemental Table 4). Among the overlapping genes of RNA-seq and ChIP-seq, Teosinte branched1/Cycloidea/Proliferating cell factor (TCP) transcription factors, including TCP18 (BRC1) were identified (Supplemental Table 3). TCP TFs have profound effects on the growth patterns of meristems and lateral organs, and *BRC1* plays a central role in shoot branching, while *BRC2* plays a slightly weaker role than *BRC1* (Aguilar-Martínez et al., 2007; Nicolas and Cubas, 2016). Axillary meristem initiation and axillary bud activation require the minimization of auxin content in leaf axil, which largely depends on the auxin efflux mediated by PIN1 (Wang et al., 2014a). And, *AXR1* is necessary for auxin to inhibit axillary bud growth (Stirnberg et al., 1999). The qRT-PCR analysis showed that *BnBRC1*, *BnBRC2*, and *BnAXR1* were down-regulated in *BnaA03.WRKY28* overexpression lines compared to those in WT, while *BnPIN1* was up-regulated (Figure 9D). In addition, two W-box elements were found in about 1,000 bp upstream of the start codon of *BnaC03.BRC1*. Sequence containing two W-box elements in the *BnaC03.BRC1* promoter was synthesized as probe. In EMSA assay, BnaA03.WRKY28 bound to DNA probe to produce a specific shifted band, which was diluted when unlabeled probe was added (Figure 9E). These results support that *BnaA03.WRKY28* is involved in growth and development, especially in axillary buds outgrowth and branches formation.

**Figure 9.**
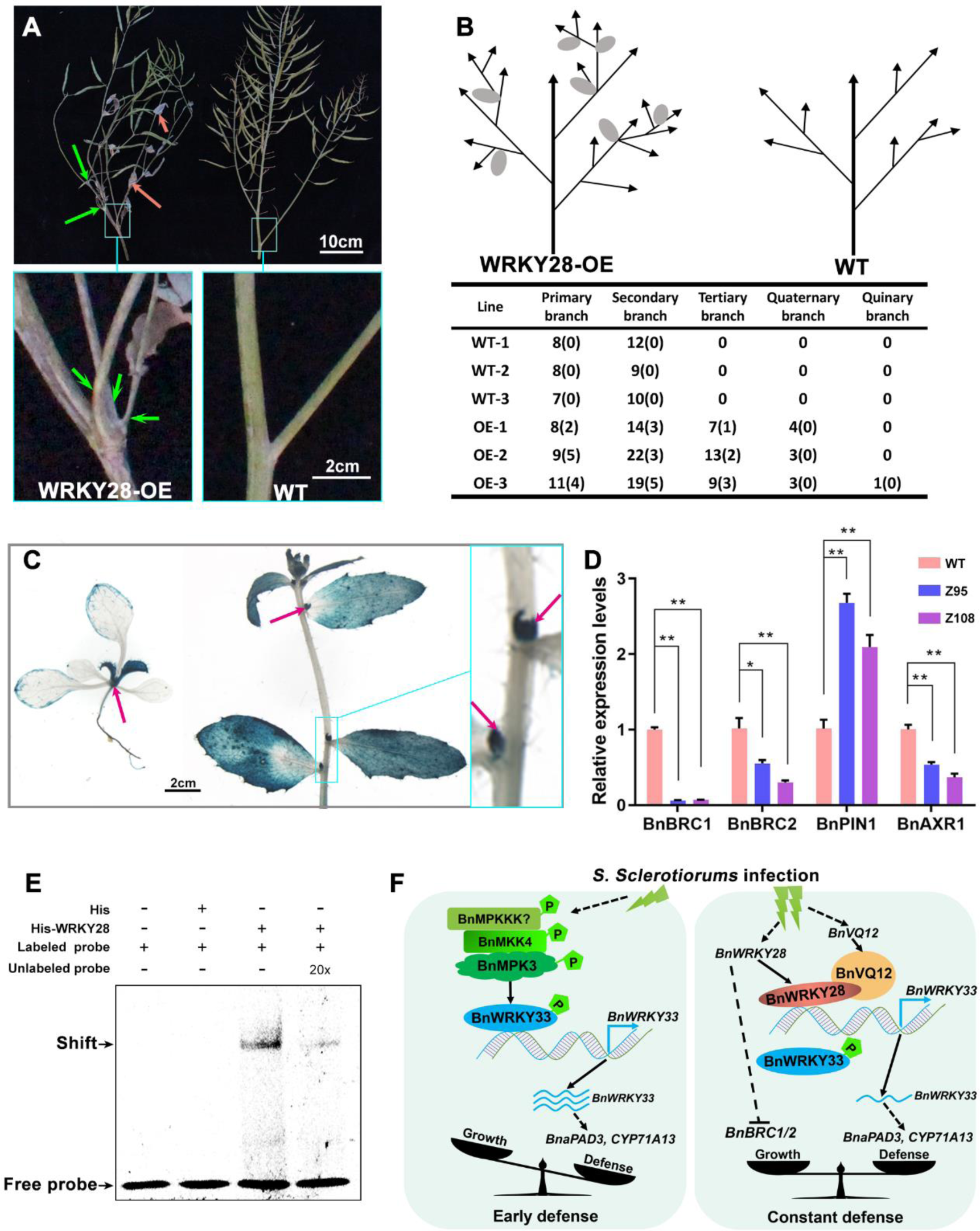
BnaA03.WRKY28 Promotes Axillary Branch Formation. **(A)**Compared with WT, *BnaA03.WRKY28* overexpression lines produced two or more branches in one axil (indicated by green arrows) and exhibited late ripening with many green cauline leaves (indicated by magenta arrows). Scale bars, 10 cm (top image), 2 cm (bottom images). Bottom, close-up view of the representative leaf axil in the square in the top image. **(B)** Schematic representation and statistics of lateral branches or buds in the leaf axils in *BnaA03.WRKY28* overexpression (WRKY28-OE) lines and WT plants in the late growth stage of WT plants (n=3). Top, schematic representation of branches in WRKY28-OE lines and WT plants. The higher- order branches and cauline leaves were common in WRKY28-OE lines. The gray ellipses indicate cauline leaves. Bottom, statistical analysis of the number of branches or buds in each order of branching and the number of branching positions that produce two or more branches in one axil (shown in brackets). The phenotype was investigated with similar results in the field for 2 years. **(C)** GUS staining showing that *BnaA03.WRKY28* was strongly expressed around the leaf axils. Left, 2-week-old plant. *BnaA03.WRKY28* was strongly expressed in the latest true leaves and the axils of rosette leaves (indicated by magenta arrows). Middle, 6-week-old plant. The expression of *BnaA03.WRKY28* could be detected in the leaf axils (arrows indicated). Right, close-up view of the leaf axils in the square in the middle image. Scale bars, 2 cm (left and middle image). **(D)** qRT-PCR for examining the transcription levels of branching related genes in *BnaA03.WRKY28* overexpression lines and WT. **(E)** EMSA to detect the possibility of BnaA03.WRKY28 binding to the *BnaC03.BRC1* promoter. The sequence containing two W-box motifs in the *BnaC03.BRC1* promoter is used as a probe. **(F)** Model showing the actions of *BnWRKY33* and *BnaA03.WRKY28* in response to *S. sclerotiorum*.

## DISCUSSION

No germplasm is immune to Sclerotinia in Brassicaceae species, which has made the breeding of resistant varieties less effective for decades, and the potential molecular mechanisms of rapeseed response to this pathogen are still poorly understood. We previously characterized the positive regulatory role of BnWRKY33 in Sclerotinia resistance (Liu et al., 2018). Here, we focused on two BnWRKY TFs (BnaA03.WRKY28 and BnWRKY33) that play an opposite role in response to Sclerotinia by competitive binding to the promoter of *BnWRKY33*. Taking our results together, we propose a hypothetical model that accounts for the actions of *BnWRKY33* and *BnaA03.WRKY28* in response to *S. sclerotiorum* (Figure 9F). When threatened by Sclerotinia, the innate immune system of rapeseed plants is activated and multiple disease resistance mechanisms will work. Among these mechanisms, the MAPK cascade signal pathway plays an important role. Soon, BnaA06.MPK3 and BnaC03.MPK3 are activated when phosphorylated by BnaA03.MKK4. Activated BnMPK3 phosphorylates BnWRKY33 and activates the transcriptional activity of BnWRKY33. BnWRKY33 binds to its own promoter to maintain the expression of *BnWRKY33* at a high abundance. BnWRKY33 regulates the high expression of disease resistance-related genes, such as *BnPAD3* and *BnCYP71A13*, which enhance the biosynthesis of phytoalexin and inhibit the invasion of the fungal pathogen. Defense is dominant during this stage. Over constant infection, the plant is at a potential risk of resistance breakdown and disease resistance continues to be highly costly. Then, *BnaA03.WRKY28* and *BnaA09.VQ12* are deeply induced. BnaA03.WRKY28 can also bind to the promoter of *BnWRKY33*, and BnaA09.VQ12 helps BnaA03.WRKY28 compete with BnWRKY33 to bind to the promoter of *BnWRKY33* by interacting with BnaA03.WRKY28. Because the transcriptional activity of BnaA03.WRKY28 is lower than that of BnWRKY33, the expression of *BnWRKY33* is lower than that mediated by BnWRKY33 during the early infection stage. So the expression of resistance genes regulated by BnWRKY33 will also decrease, which is equivalent to the braking effect of disease resistance. Furthermore, *BnaA03.WRKY28,* which is expressed in leaf axils, stimulates the initiation of axillary meristem and the development of axillary buds that grow into complete branches. At this stage, the resistance strength decreases and the growth returns to be active. This brake of disease resistance contributes to the trade-off between defense and growth, and may provide the possibility for the plants to ensure their survival.

Disease resistance is essential to ensure the continuation of life when threatened by pathogens; however, disease resistance is costly. The immune response leads to the inhibition of plant growth and development in Arabidopsis (Gómez - Gómez et al., 1999; Tian et al., 2003). In rice, *OsWRKY45* plays a key role in SA signaling pathway and mediates resistance to broad pathogens; however, overexpression of *OsWRKY45* to enhance disease resistance is accompanied by poor agronomic traits (Goto et al., 2015). Disease resistance, especially constitutive defense reactions with overexpression of resistance-related signaling molecules and TFs are associated with side effects on agronomic traits, such as lesion mimic phenotypes, reduced height, and early senescence (Delteil et al., 2010). In this study, we found that constitutively active BnaA03.MKK4^DD^, but not BnaA03.MKK4, induced hypersensitive response cell death (Supplemental Figure 10). A trade-off between plant growth and disease resistance has become a generally accepted concept (Karasov et al., 2017). Allocation of limited resources to various life activities may be responsible for the trade-off between growth and defense (Heil and Baldwin, 2002; Matyssek et al., 2005). Phytoalexins, a class of low molecular mass antimicrobial compounds synthesized *de novo* and accumulated only after the plant is infected, are significant component in pathogen defense (VanEtten et al., 1994; Ahuja et al., 2012). Camalexin, the major phytoalexin in Arabidopsis and some Brassicaceae species, plays a role in defense against *S. sclerotiorum* and *B. cinerea* (Mao et al., 2011; Stotz et al., 2011; Ahuja et al., 2012). The biosynthesis of camalexin is derived from tryptophan, which is also the precursor substance of IAA (indole-3-acetic acid) (Cohen et al., 2003; Glawischnig et al., 2004). This may contribute to the antagonism between growth and defense. Two P450 enzymes, CYP71A13 and PAD3, are involved in camalexin biosynthesis. In the absence of stress, camalexin biosynthesis-related genes, including *AtCYP71A13* and *AtPAD3*, are expressed at very low levels, but are highly induced when targeted by WRKY33 under pathogen infection to activate camalexin biosynthesis (Nafisi et al., 2007; Qiu et al., 2008; Zhou et al., 2020). Our previous study demonstrate that the transcript abundance of *BnCYP71A13* and *BnPAD3* is significantly increased in *BnWRKY33* overexpression lines (Liu et al., 2018). In this study, we found that *BnMPK3/BnaA03.MKK4* overexpression lines, and *BnaA03.WRKY28*/*BnaA09.VQ12* knockout lines exhibited better disease resistance with higher expression levels of *BnCYP71A13* and *BnPAD3* compared with wild-type plants.

*BnaA03.WRKY28* overexpression lines showed more active axillary meristem and produced more branches or buds in the leaf axils. In Arabidopsis, *AtWRKY28* and *AtWRKY71* are in the same clade, and overexpression of *AtWRKY28* and *AtWRKY71* both cause excessive branching (Guo et al., 2015). *AtWRKY71* has been demonstrated to promote axillary meristem initiation by positively regulating the key branch meristem formation-related TFs, including AtRAX1, AtRAX2, and AtRAX3 (Guo et al., 2015). In our RNA-seq and ChIP-seq data, lateral branch formation-related genes, such as TCP TFs (including BnBRC1) and auxin polar transporter (including BnPIN1), were identified. The changes in expression level of these genes effectively resulting in excessive branching of *BnaA03.WRKY28* overexpression lines.

Since VQ proteins were identified in plants, most VQ proteins are thought to regulate genes expression by interacting with WRKY TFs to be involve in plant growth, development and stress response (Cheng et al., 2012; Jing and Lin, 2015). In Arabidopsis, the WRKY superfamily TFs contain 74 members, which are classified into three groups (group I, group II, and group III) (Eulgem et al., 2000), and the group II is further divided into IIa, IIb, IIc, IId, and IIe (Rushton et al., 2010). Most VQ proteins physically interact with group I or group IIc WRKY proteins through their WRKY domain (Cheng et al., 2012; Jing and Lin, 2015). Out of the 74 members of the WRKY, there are 13 members of group I and 19 members of group IIc. This total number (32) is similar to the total number (34) of VQ proteins in Arabidopsis (Cheng et al., 2012). The interaction between VQ and WRKY can affect the DNA-binding activity of WRKY TFs (Jing and Lin, 2015). WRKY8 can interact with VQ9 and VQ10, respectively. VQ9 can weaken the DNA binding activity of WRKY8, but VQ10 can enhance the binding ability of WRKY8 to downstream target genes. Here, we identified another VQ protein, BnaA09.VQ12, which interacted with BnaA03.WRKY28 by binding to DNA binding domain of BnaA03.WRKY28. Formation of the BnaA09.VQ12-BnaA03.WRKY28 complex enhances the DNA binding activity of BnaA03.WRKY28.

Plants have evolved intricate and precise gene regulatory networks to modulate their own growth and cope with the changing environment. It is well established that regulatory proteins usually play a synergistic or antagonistic role in biological functions. *AtWRKY33* positively contributes to salt tolerance, *Alternaria brassicicola* and *Botrytis cinerea* resistance (Jiang and Deyholos, 2009; Birkenbihl et al., 2012; Liu et al., 2017). *BnWRKY33* (homologous gene of *AtWRKY33*) plays a positive role in Sclerotinia resistance (Wang et al., 2014b; Liu et al., 2018). WRKY33 may be a key TF in the plant stress response. In Arabidopsis, MEKK1-MKK4/5-MPK3/6 module is well studied and acts as a positive regulator of defense responses (Ichimura et al., 2002; Meng and Zhang, 2013). Next, MPK3/6 plays roles in defense through phosphorylation of their substrates, such as WRKY33 and ERF104 (Bethke et al., 2009; Mao et al., 2011). In this study, we verified that phosphorylation and activation of BnWRKY33 by the BnMKK4-BnMPK3 module is necessary for a transcriptional burst of *BnWRKY33*. The W-box is the target site of the WRKY superfamily TFs. Both the promoters of *AtWRKY33* and *BnWRKY33* contain a set of three W-boxes, and WRKY33 can directly bind to its own promoter (Mao et al., 2011; Liu et al., 2018). Additionally, BnWRKY15 can also target the promoter of *BnWRKY33* (Liu et al., 2018). In this study, we identified another WRKY TF (BnaA03.WRKY28) that can bind to the promoter of *BnWRKY33*. Specifically, BnWRKY33 and BnWRKY15 bind to the W1 of the *BnWRKY33* promoter (Liu et al., 2018), whereas BnaA03.WRKY28 can target W1 and W3. In addition, *BnWRKY33* and *BnaA03.WRKY28* exhibited different induced trends within 0-72 h after infection. *BnWRKY33* was significantly induced earlier, peaked at 24 h, and then decreased, while *BnaA03.WRKY28* was induced later and reached its peak at 48 h. The decreased expression level of *BnWRKY33* was accompanied by increased expression level of *BnaA03.WRKY28*, which was caused by the state when lower-transcriptional-activity BnaA03.WRKY28 replaced higher-transcriptional-activity BnWRKY33 and preferentially bound to the promoter of *BnWRKY33* in the late stage of infection. BnaA03.WRKY28 and BnWRKY33 were competitive when binds to the *BnWRKY33* promoter. Competitive binding is commonly reported in protein-protein interactions. SAUR17 and SAUR50 can interact with PP2C-D1, but SAUR17 has a higher affinity for PP2C-D1. In the dark, PP2C-D1 preferentially interacts with SAUR17 to promote apical hook development (Wang et al., 2020). In root stele cells, abundant accumulation of FIT-binding protein (FBP) takes precedence over bHLH038/039/100/101 in interactions with FIT to maintain

Fe and Zn homeostasis in plants (Chen et al., 2018a). From this study, it is suggested that BnaA03.WRKY28 has a higher affinity for the *BnWRKY33* promoter than BnWRKY33 with the help of BnaA09.VQ12. This counts much for fine-tuning of defense strength.

## MATERIALS & METHODS

### Plant Materials and Growth Conditions

All transgenic lines in this study were established from the spring varieties Jia 9709 and Westar background. Transgenic lines developed in this study include *BnaA03.WRKY28-OE*/Jia 9709 and Westar, *BnaA03.WRKY28* CRISPR/Cas9-mediated mutant (*BnaA03.WRKY28-CR*/ Jia 9709), *BnaA09.VQ12-OE*/Jia 9709, *BnaVQ12* CRISPR/Cas9-mediated mutant (*BnaVQ12-CR*/Jia 9709), *BnaA06.MPK3-OE*/Jia 9709 and Westar, *BnaC03.MPK3-OE*/Jia 9709 and Westar, *BnMPK3* CRISPR/Cas9-mediated mutant (*BnMPK3-CR*/Jia 9709 and Westar), and *BnaA03.MKK4-OE*/Jia 9709. Plants were grown at the experimental station in Wuhan under natural conditions for stem inoculation and seed propagation. For rapeseed leaf inoculation and Arabidopsis or tobacco growth, seedlings were grown in greenhouse at 22°C under long-day conditions (16 h day/8 h night).

### Plasmid Construction and Plant Transformation

To generate the overexpression construct, the full-length coding sequences of *BnaA03.WRKY28*, *BnaA09.VQ12*, and *BnaA03.MKK4* were cloned into the plant binary vector pCAMBIA2300 under the control of CaMV35S, in which BnaA03.WRKY28 and BnaA03.MKK4 were fused with FLAG tag. The CDS of two copies of BnMPK3 was inserted into the pCAMBIA1300-CaMV35S-mcherry vector. The gene editing constructs for *BnaA03.WRKY28*, *BnaVQ12*, and *BnaMPK3* via CRISPR/Cas9 were designed using the method described by Chen’s lab using the binary vector *pKSE401* (Xing et al., 2014). The sgRNA of each gene was designed using CRISPR-P (http://cbi.hzau.edu.cn/cgi-bin/CRISPR). All constructs were confirmed by sequencing. All of the recombinant plasmids were introduced into *Agrobacterium* strain GV3101 and transferred to different genetic backgrounds, according to the transformation method as previously described (Zhang et al., 2020a). Primers for constructs are listed in Supplemental Table 5.

### Pathogen Inoculation and Disease Resistance Assay

*S. sclerotiorum*, A367, kindly provided by Prof. Daohong Jiang from Huazhong Agriculture University (Wuhan, China), was used for inoculation. The *S. sclerotiorum* isolate was cultured on potato dextrose agar medium (20% potato, 2% dextrose, and 1.5% agar) at 22°C in the dark. For leaf inoculation, the assay was performed according to previous study (Liu et al., 2018; Wang et al., 2018). The latest fully unfolded leaves of rapeseed in approximately 4 weeks were removed from plants, and the detached leaves were placed on soaked gauze. Then the mycelial side of the fungus agar block (7 mm in diameter) was attached to the leaf surface. The inoculated leaves were covered with plastic films to maintain moisture at 22°C. At 48 h post-inoculation, images were taken of inoculation leaves and lesion areas were measured using ImageJ software. For stem inoculation, the assay was performed as previously described (Wei et al., 2016). At the end of flowering time, the mycelial side of the fungus agar block (7 mm in diameter) was attached to the stem, which was approximately 30 cm above the soil with plastic wrap. The stems were sprayed with water mist. After 96 h inoculation, images were taken of inoculation stems and lesion length was measured using ImageJ software.

### RNA Extraction and qRT-PCR

Total RNA was extracted from leaves of transgenic lines and wild-type plants treated with pathogens or hormones using the RNAprep pure plant kit (TIANGEN, DP441, Beijing, China). Then, 1 µg of total RNA was used for cDNA synthesis using the RevertAid™ First Strand cDNA Synthesis Kit (Fermentas, #K1622, USA). qRT-PCR analysis was performed with the CFX96^TM^ Real-Time system (Bio-Rad, USA) using the SYBR Green Realtime PCR Master Mix (TOYOBO, QPK-201, Japan). BnACTIN2 was used as a control to normalize expression levels according to the 2^-ΔΔCT^ method (Livak and Schmittgen, 2001). Primers used in the qPCR assays are given in Supplemental Table 5.

### Yeast One-Hybrid Assay

To verity the binding of BnaA03.WRKY28 and *BnWRKR33* promoter, yeast one-hybrid assays were performed as the manual of Matchmaker Gold Yeast One-Hybrid System (Clontech). The coding sequence of BnaA03.WRKY28 was cloned into pGADT7 vector. The bait plasmid containing the promoter fragment was constructed as previous study (Liu et al., 2018), and was transformed into the Y1HGold yeast strain. Then the prey plasmid was introduced into the bait strain. Clones were cultured on selective dropout medium SD/-Leu with or without 300 ng/mL AbA.

### Yeast Two-Hybrid Assay

The full-length CDS of BnWRKY33, BnaA03.MKK4, BnaA09.VQ12, and BnaA01.VQ22 were cloned into the pGADT7 vector, and the CDS of BnaA03.WRKY28, truncated BnaA03.WRKY28, BnaA06.MPK3, and BnaC03.MPK3 were cloned into the pGBKT7 vector. To assay protein-protein interactions, the recombinant pGADT7 and pGBKT7 constructs were cotransformed into AH109 yeast strain. The Y2H assays were performed following the manufacturer’s protocol using the Matchmaker GAL4 Two-Hybrid System (Clontech). Clones were grown on selective dropout medium (lacking either Trp and Leu or Trp, Leu, His and Ade).

### Protein Expression and Purification

BnaA03.WRKY28, BnaA09.VQ12, BnWRKY33, BnWRKY33-N, BnWRKY33^AA^-N, BnaA03.MKK4, and BnaA03.MKK4^DD^ were cloned into pET32a to generate His-fusion proteins, and BnaA03.WRKY28, BnaA03.WRKY28 164-227, BnaA06.MPK3, and BnaC03.MPK3 were inserted into pGEX4T-1 to generate GST-fusion proteins, and BnWRKY33 was also fused with MBP tag in the pMAL-c2x vector. Except for the constructs that containing full-length CDS of BnWRKY33, which were transformed into *E. coli* Rosetta (DE3), fusion proteins were induced using *E. coli* BL21. The expression of fusion proteins was induced by 0.2-0.5mM IPTG at 16°C. After 16 h of induction, the cells were collected and resuscitated with the corresponding lysis buffer. The cells were lysed using a high-pressure cell disrupter (JNBIO, China), and the supernatant was added onto a column equipped with Ni^2+^ affinity resin (BBI, C600033, Shanghai, China),

GST-Sefinose resin (BBI, C600031, Shanghai, China), or amylose resin (Sangon, C500096, Shanghai, China) after centrifugation at 12000g at 4°C for 30min. The fused proteins were washed five times with the corresponding buffer, and the His-fused proteins were eluted with a buffer containing 250mM imidazole, the GST-fused proteins were eluted with 0.5 mM reduced glutathione, and the MBP-fused proteins were eluted with 10 mM maltose. The purified protein was examined using sodium dodecyl sulfate polyacrylamide gel electrophoresis (SDS-PAGE) and visualized with Coomassie blue staining.

### Electrophoretic Mobility Shift Assay (EMSA)

For EMSA, His-tagged BnaA03.WRKY28 protein was used. Two complementary oligonucleotide strands isolated from the promoter of *BnWRKY33* were labeled with Cy5 and annealed to generate probes. The DNA-protein binding activities were incubated in the reaction system containing EMSA/Gel-shift Binding Buffer (Beyotime, GS005, China) and 25 nM Cy5-labeled probe at 23°C for 30 min. For competition, 50-fold unlabeled probes were added to the reactions. The reaction mixture was loaded on a 6% native polyacrylamide gel and electrophoresed in 0.5 × TBE (45 mM Tris-base, 45 mM boric acid, 0.5 mM EDTA, pH 8.3) at 4°C for 1 h at 80 V in the dark. Fluorescence-labeled DNA on the gel was directly detected using Fujifilm FLA-9000 (FujiFilm, Japan).

### Chromatin Immunoprecipitation Assay

The methods used for Chromatin Immunoprecipitation (ChIP) were previously described (Zhang et al., 2020b). Briefly, rapeseed leaves were crosslinked in 1% formaldehyde for 15 min and quenched with 0.2 M glycine. Samples were lysed in buffer (50 mM HEPES-KOH pH7.5, 150 mM NaCl, 1 mM EDTA, 0.1% sodium deoxycholate, 1% Triton X-100, 1% SDS) for 10 min at 4°C, and the chromatin was fragmented by sonication using a Bioruptor (Diagenode). The chromation solution was incubated with 10 ul anti-FLAG (Sigma) for 6 h at 4°C. The immunoprecipitated chromatin was washed subsequently with low-salt buffer (50 mM HEPES-KOH, 150 mM NaCl, 1 mM EDTA, 0.1% sodium deoxycholate, 1% Triton X-100, 0.1% SDS), high-salt buffer (50 mM HEPES-KOH, 350 mM NaCl, 1 mM EDTA, 0.1% sodium deoxycholate, 1% Triton X-100, 0.1% SDS), wash buffer (10 mM Tris-HCl pH 8.0, 250 mM LiCl, 0.1% sodium deoxycholate, 0.5% NP-40, 1 mM EDTA), and TE buffer (10 mM Tris-HCl pH 8.0, 1 mM EDTA). The protein-DNA complexes were eluted by Elution buffer (50 mM Tris-HCl pH 7.5, 10 mM EDTA, 1% SDS) for 15 min at 65°C with agitation at 900 rpm. ChIP DNA was extracted with phenol:chloroform:isoamyl alcohol (Sigma-Aldrich, P3803), precipitated with ethanol, and resuspended in TE buffer, and used for ChIP-qPCR or sequencing (ChIP-seq). The primers used for ChIP-qPCR are listed in Supplemental Table 5.

### Arabidopsis Protoplast Isolation and Transient Assay

In this study, Arabidopsis protoplast isolation and transient transformation were widely used, such as subcellular localization, bimolecular fluorescence complementarity and dual-luciferase test. Arabidopsis protoplast transient expression assays were performed according to previously described protocols (Yoo et al., 2007). In brief, the 5th-7th true leaves of healthy 4-week-old Arabidopsis plants were cut into 0.5-1mm wide filaments. The cut leaves were hydrolyzed in the enzymatic hydrolysate (1:1.5% CellulaseR10, 0.2% 0.4% MacerozymeR10, 0.4 M Mannitol, 20 MMMES (pH5.7)) for 3 h in the dark, and the reaction was terminated with the same volume of W5 (154 mm NaCl, 125 mM CaCl_2_, 5 mM KCl_2_, 2 mM MES (pH5.7)). After centrifugation at 100g for 3 min at 4°C, the protoplasts were re-suspended with W5 and placed on ice for 30 min. Then, the protoplasts were centrifuged under the same conditions and re-suspended with MMG (0.2 M Mannitol, 0.2 mm MgCl_2_, 4 mm MES (pH5.7)). Protoplasts were transformed mediated by PEG. The total 20 μg plasmids were incubated with 200 μL protoplasts and 20% PEG solution for 10 min at room temperature and then quenched with W5. The supernatant was removed by centrifugation at room temperature and cultured with 400 μL WI (4 mM MES (pH5.7), 20 mM KCl, 0.5M Mannitol) for 12-18h, and then the follow-up experiments were conducted.

### Dual-luciferase Transient Transcriptional Activity Assay

Genes encoding TFs, MPK cascade, and VQ12 examined in this study were cloned into pGreenII 62-SK and pGAL4BD vectors as effectors (Hellens et al., 2005; Zong et al., 2016). Promoter sequences of *BnWRKY33* were inserted into pGreenII 0800-LUC vector to construct reporters. CaMV35S-driven *Renilla* LUC was used as an internal control. The effector, reporter, and internal control were cotransformed into Arabidopsis protoplasts, and LUC activity was measured according to the manufacturer’s instructions (Promega, E1910, USA).

### *In Vitro* Pull-Down Assay

For *in vitro* pull-down assays, His-tagged BnWRKY33, BnaA03.MKK4, and BnaA09.VQ12, and GST-tagged BnaA06.MPK3, BnaC03.MPK3, BnaA03.WRKY28, and BnaA03.WRKY28 164-227 proteins were used. GST or GST-tagged protein sample was incubated with GST beads at 4°C for 1h, followed by washing five times, and incubation with His-tagged protein at 4°C for 2 h. The beads were then washed five times. The isolated precipitate was boiled with loading buffer at 100°C for 6 min and detected by western blot using anti-His (Abclonal, AE003, China, 1:10000) or anti-GST (Abclonal, AE001, China, 1:10000) antibodies. The ECL western specific luminescence detection kit (BIO-RAD, #1705060, USA) was used to collect the signal, and the images were scanned using the Image Quant luminescence LAS 4000 machine.

### Bimolecular Fluorescence Complementation (BiFC) Assay

The BiFC vectors used in this study were based on the research of Waadt et al. (Waadt et al., 2008). BnaA03.WRKY28 and BnaA03.WRKY28 164-227 were amplified and cloned into the pSPYCE vector, and the full-length CDS of BnaA09.VQ12 and BnaA01.VQ22 were inserted into the pSPYNE vector. The constructs were transformed into Arabidopsis protoplasts in different combinations. After 16 h incubation in the dark, the fluorescence signals were visualized under a laser scanning confocal microscope (TCS SP2; Leica).

### CoIP Assay

BnaA06.MPK3, BnaC03.MPK3, BnaA03.WRKY28, and BnaA03.WRKY28 164-227 were fused with GFP tag. BnWRKY33, BnaA03.MKK4, and BnaA09.VQ12 were fused with FLAG tag. Four-week-old *Nicotiana benthamiana* leaves were infiltrated with GV3101 *Agrobacterium* strain carrying constructs combination. After incubation for 3 days, the total protein was extracted from 5 g infiltrated leaves with 5 mL extraction buffer (50 mM Tris-HCl pH 7.5, 150 mM NaCl, 1% Triton X-100, 5 mM EDTA, 10% glycerol, and 1 × protease inhibitor (Roche)) and incubated with 25 μL GFP-Trap®-MA (Chromotek, gtma-20, Germany) at 4°C for 1 h. Then, the beads were washed five times with wash buffer (50 mM Tris-HCl pH 7.5, 150 mM NaCl, 0.1% Triton X-100, 5 mM EDTA, and 10% glycerol) and eluted with 50 μL SDS loading buffer. The immunoblot assays were conducted with anti-FLAG (Abclonal, AE005, China, 1:10000) and anti-GFP (Abclonal, AE012, China, 1:10000) antibodies. The ECL western specific luminescence detection kit (BIO-RAD, #1705060, USA) was used to collect the signal, and the images were scanned using the Image Quant luminescence LAS 4000 machine.

### *In Vitro* Phosphorylation Assay

The phosphorylation assays were performed as previously described (Mao et al., 2011). Briefly, 0.5 μg GST-tagged BnaA06.MPK3/BnaC03.MPK3 was activated by 0.1 μg His-tagged BnaA03.MKK4^DD^ in the reaction buffer (50 mM Tris-HCl pH 7.5, 50 μM ATP, 10 mM MgCl_2_ and 1 mM DTT) at 25°C for 1 h. Then, activated BnaA06.MPK3/BnaC03.MPK3 was used to phosphorylate BnWRKY33 and BnWRKY33-N (5:1 substrate/enzyme ratio) in the reaction buffer (50 mM Tris-HCl pH 7.5, 10 μM ATP, 10 mM MgCl_2_, 1 μCi [γ-^32^P] ATP and 1 mM DTT) at 25°C for 1 h. The reactions were stopped with the SDS loading buffer. Phosphorylation signals for BnWRKY33 and BnWRKY33-N were visualized by autoradiography after being resolved in a 10% SDS-PAGE gel.

### Subcellular Localization

In order to determine the subcellular localization of BnaA03.WRKY28 and BnaA09.VQ12, full-length CDS without stop codon was cloned into pM999 vector to construct a recombinant of fused green fluorescent protein driven by CaMV35S. The OsGHD7 fusion CFP or RFP located in the nucleus was used as the control. The fusion constructs were transformed into Arabidopsis protoplasts. After incubation in the dark for 16 h, the fluorescence signals were visualized under a laser scanning confocal microscope (TCS SP2; Leica).

### Histochemical GUS Staining

The samples of *ProBnaA03.WRKY28:GUS* transgenic plants were immersed in 90% acetone on ice for 20-30 min, then washed twice with 50 mM phosphate buffer (mixing 39 mL 0.2M NaH_2_PO_4_ꞏ2H_2_O and 61 mL 0.2M Na_2_HPO_4_ꞏ12H_2_O in 400 mL solution, pH 7.2). Thereafter, the samples were stained in 1 mg/mL X-Gluc solution overnight at 37 °C, and 70% ethanol was used to remove the chlorophyll. Images were captured using a Leica stereoscope.

### Accession Numbers

The accession numbers from http://www.genoscope.cns.fr/colza-ggb/ are as follows: *BnaA03.WRKY28* (BnaA03g43640D), *BnWRKY33* (BnaA05g34850D), *BnaA09.VQ12* (BnaA09g42280D), *BnaA06.MPK3* (BnaA06g18440D), *BnaC03.MPK3* (BnaC03g55440D), *BnaA03.MKK4* (BnaA03g36120D), *BnaC03.BRC1* (BnaCnng23770D).

## Supplemental Data

**Supplemental Figure 1.** Sequences analysis of *BnWRKY28*.

**Supplemental Figure 2.** The phenotypic investigation of *BnaA03.WRKY28* transgenic lines on Sclerotinia resistance.

**Supplemental Figure 3.** ChIP-seq and RNA-seq analysis of *BnaA03.WRKY28* overexpression lines.

**Supplemental Figure 4.** BnaA03.WRKY28 binds to W1/W3 of *BnWRKY33* promoter in yeast.

**Supplemental Figure 5.** Expression profiling analysis of the MAPK cascade in response to Sclerotinia infection.

**Supplemental Figure 6.** BnaA06.MPK3/BnaC03.MPK3 interacts with BnaA03.MKK4 and BnWRKY33 *in vitro*.

**Supplemental Figure 7.** B*n*aA03*.MKK4* positively contributes to *S. sclerotiorum* resistance in *Brassica napus*.

**Supplemental Figure 8.** Subcellular localization of BnaA03.WRKY28 and BnaA09.VQ12 in Arabidopsis protoplasts.

**Supplemental Figure 9.** B*n*aA03*.WRKY28* overexpression lines exhibit excessive branches.

**Supplemental Figure 10.** Constitutively active BnaA03.MKK4^DD^ induce hypersensitive response cell death.

**Supplemental Table 1.** Peaks identified in ChIP-seq.

**Supplemental Table 2.** List of BnaA03.WRKY28-regulated genes in Brassica napus.

**Supplemental Table 3.** List of overlapping genes bound and regulated by BnaA03.WRKY28.

**Supplemental Table 4.** Genes used for heatmapping.

**Supplemental Table 5.** All primers used in this study.

## ACKNOWLEDGMENTS

We thank Prof. Daohong Jiang (Huazhong Agriculture University) for providing *S. sclerotiorum* strains, Prof. Xingwang Li (Huazhong Agriculture University) for ChIP assays assistance, Yongliang Wang for phosphorylation assays assistance, Pugang Yu for ChIP-seq and RNA-seq data analysis, and Dr. Qiang Xin and Bao Yang for manuscript modifying. This work was financed by the funding from the National Key Research and Development P rogram of China (2016YFD0100305), and the National Natural Science Foundation of China (31376120).

## AUTHOR CONTRIBUTIONS

J.T. directed the project. K.Z. and J.T. conceived and designed the research. K.Z., F.L., C.Z., and X.L. performed the experiments. K.Z., Z.W., and K.H. analyzed the data. All authors interpreted and discussed the results. K.Z. and Z.W. wrote the manuscript. K.Z., F.L., Z.W., and J.T. edited and modified the manuscript. J.W., B.Y., J.S., C.M., and T.F. supervised the research.

## Supplemental Figures

**Supplemental Figure 1.**
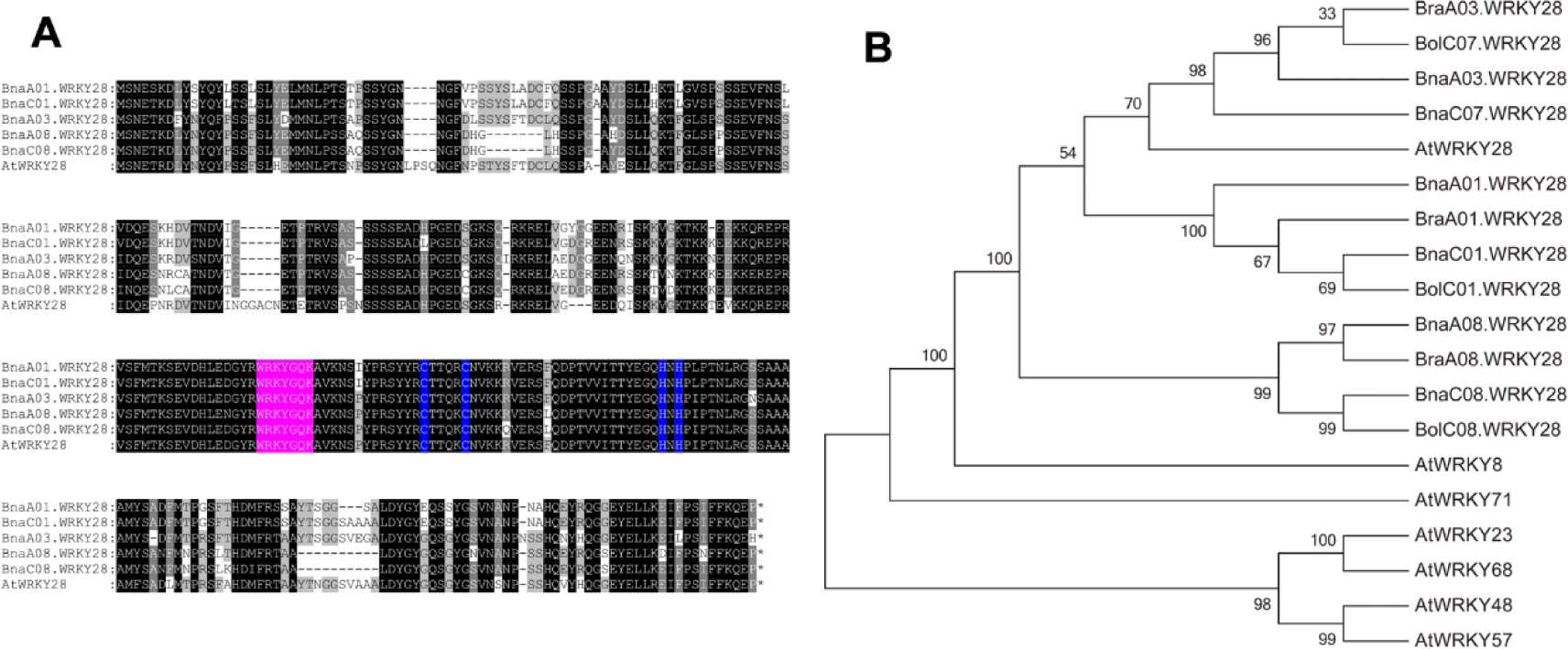
Sequences analysis of *BnWRKY28*. (Supports Figure 1) (A) Sequences of rapeseed and *Arabidopsis* WRKY28 proteins were alignmented using MEGA. Conserved residues were shown in white text on black. WRKYGQK motifs were marked in magenta, and zinc-finger motifs were marked in blue. (B) Phylogenetic analysis showing the relationship between each copy of *BnWRKY28* and *WRKY28* in *Brassica* species, as well as members of WRKY family in *Arabidopsis*. Bra, *Brassica rapa*; Bol, *Brassica oleracea*; Bna, *Brassica napus*; At, *Arabidopsis thaliana*.

**Supplemental Figure 2.**
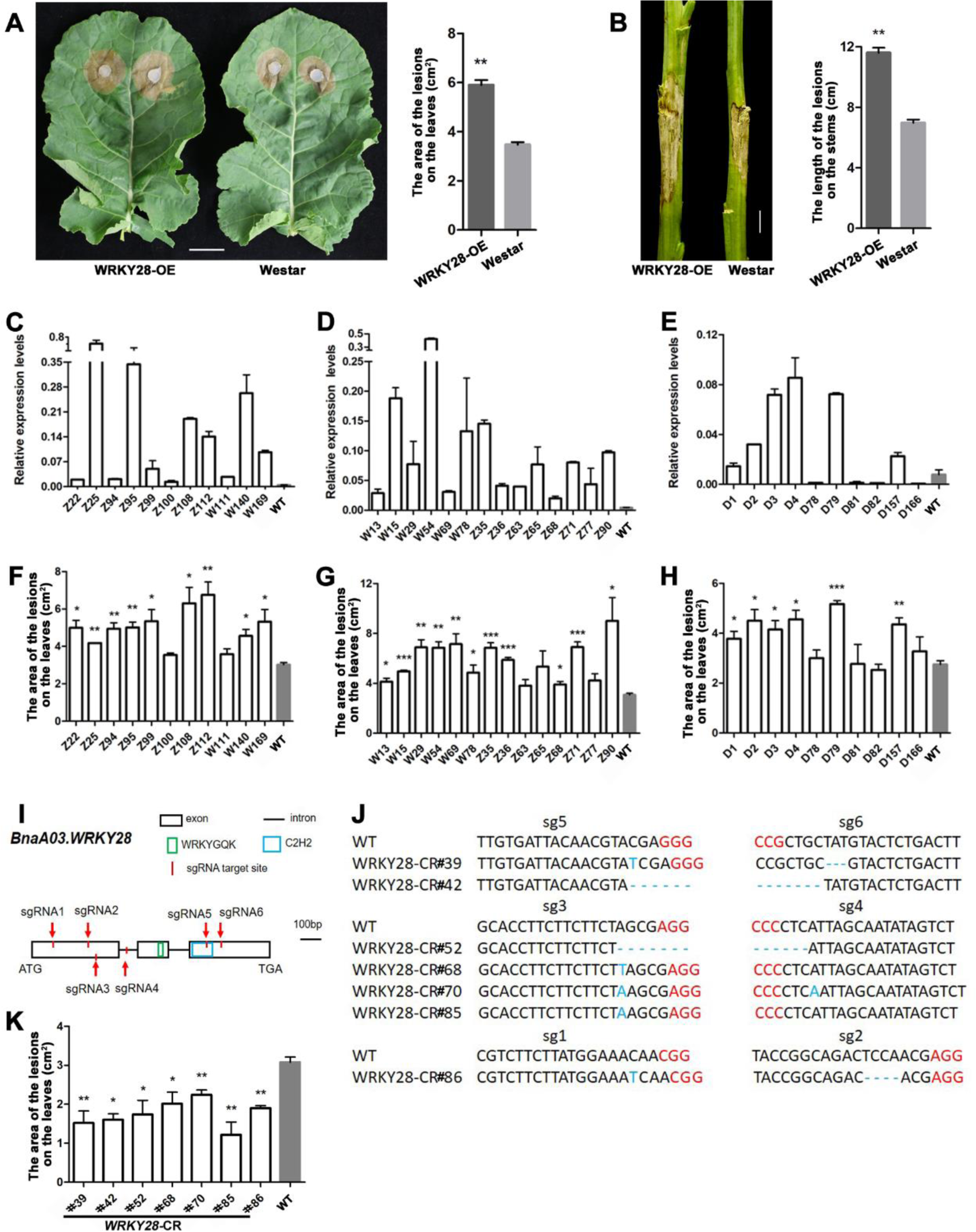
The phenotypic investigation of BnaA03.WRKY28 transgenic lines on *Sclerotinia* resistance. (Supports Figure 1) **(A)** The lesions of detached leaves from *BnaA03.WRKY28* overexpression (WRKY28-OE) lines (Westar background) and wild-type (WT) plants (Westar) in approximately 4 weeks were imaged and lesion areas were calculated when inoculated with 7 mm *S. sclerotiorum* hyphae agar block at 22°C for 48h. Scale bar, 2 cm. Data are shown as means ± SD (*n* = 3). Asterisks indicate significant differences compared with WT (t-test, \*\**P* < 0.01). **(B)** The lesion lengths on the stems of WRKY28-OE lines (Westar background) and WT plants (Westar) at the flowering stage were imaged and calculated when inoculated with *S. sclerotiorum* for 4 days. Scale bar, 2 cm. Data are shown as means ± SD (*n* = 3). Asterisks indicate significant differences compared with WT (t-test, \*\**P* < 0.01). **(C)** to **(H)** Detection of relative expression level of *BnaA03.WRKY28* in *BnaA03.WRKY28* overexpression lines and calculation of lesion areas of these lines after 48 h inoculation. **(C)** and **(F)**, Lines transformed with *BnaA03.WRKY28* fusion FLAG tag driven by CaMV35S vector in Jia9709 background; **(D)** and **(G)**, Lines transformed with *BnaA03.WRKY28* driven by CaMV35S vector in Jia9709 background; **(E)** and **(H)**, Lines transformed with *BnaA03.WRKY28* driven by CaMV35S vector in Westar background. Data are shown as means ± SD (*n* = 3). Asterisks indicate significant differences compared with corresponding WT (t-test, \**P* < 0.05, \*\**P* < 0.01). **(I)** Schematic diagram to show the characterization of the *BnaA03.WRKY28* sequence and the target sites of sgRNAs using the CRISPR/Cas9 system. **(J)** The edit types of independent homozygous mutants are shown. The protospacer-adjacent motifs (PAM) are marked in red. **(K)** Statistical analysis of lesion areas of *bnaa03.wrky28* mutants mediated by CRISPR/Cas9. Data are shown as means ± SD (*n* = 3). Asterisks indicate significant differences compared with WT (t-test, \**P* < 0.05, \*\**P* < 0.01).

**Supplemental Figure 3.**
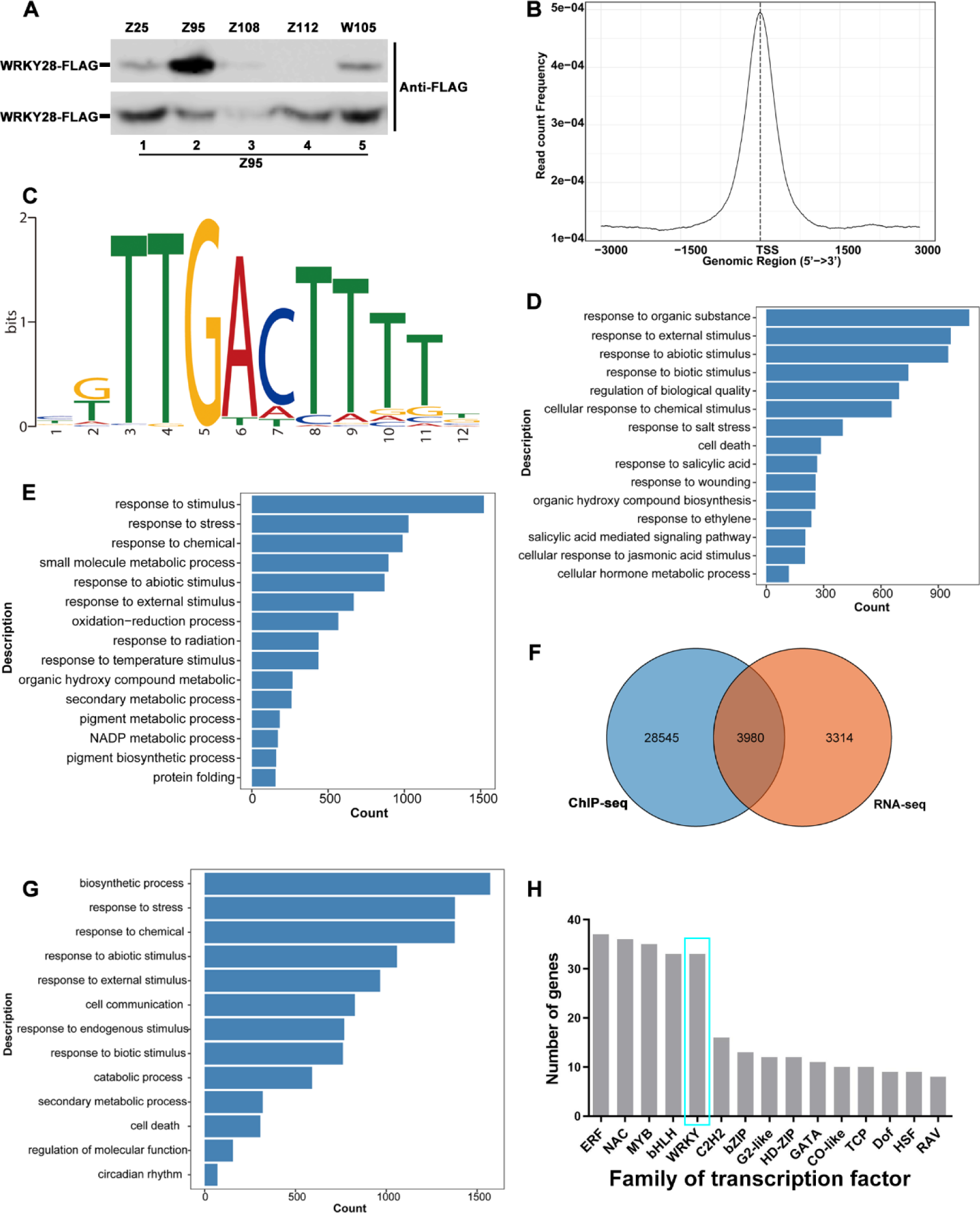
ChIP-seq and RNA-seq analysis of *BnaA03.WRKY28* overexpression lines. (Supports Figure 2) **(A)** Western blot analysis showing BnaA03.WRKY28-FLAG could be stably expressed in line Z95. Bottom image indicating the progeny plants of Z95. **(B)** Graph showing peak regions of BnaA03.WRKY28-bound associated with the transcription start site (TSS) of a gene. **(C)** MEME analysis indicates that BnaA03.WRKY28 predominantly binds to the TTGACT motif. **(D)** and **(E)** Gene ontology (GO) analysis showing significant enrichment terms were related to stimulus response in up-regulated gene cluster **(C)** and down- regulated gene cluster **(D)**. The DEGs were obtained from the comparison between line Z95 and WT. **(F)** Venn diagram showing the total number of overlapping genes identified by ChIP-seq and RNA-seq. **(G)** GO analysis shows that significant enrichment terms are related to stress response in overlapping gene cluster. **(H)** Distribution of transcription factors in overlapping gene cluster.

**Supplemental Figure 4.**
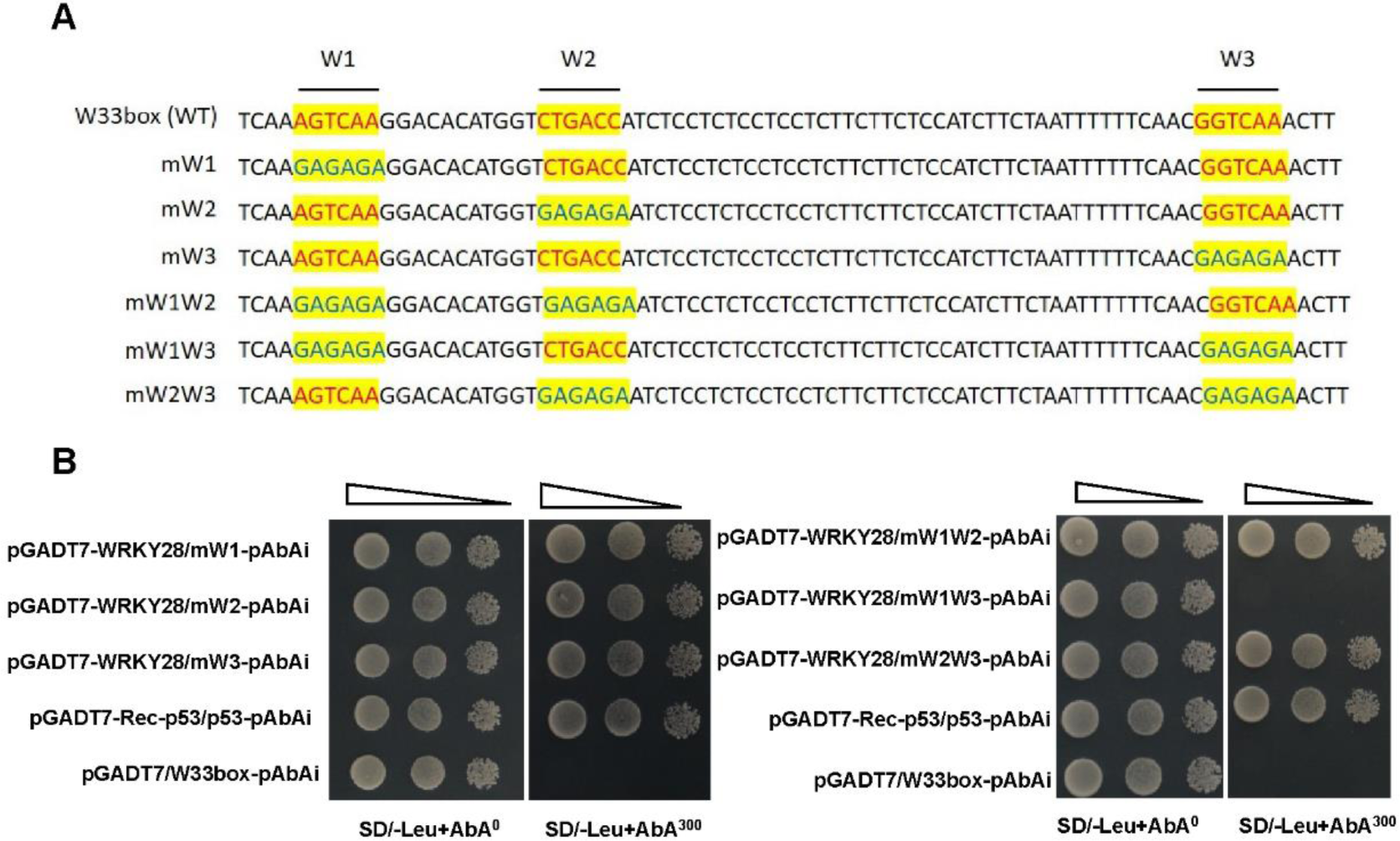
BnaA03.WRKY28 binds to W1/W3 of *BnWRKY33* promoter in yeast. (Supports Figure 2) **(A)** Schematic of wild-type and different mutation forms of W-box region of *BnWRKY33* promoter. Three W-boxes of the *BnWRKY33* promoter were mutated one by one. The core W-box sequence were mutated to GAGAGA. **(B)** Yeast one-hybrid assays show that W1 and W3, but not W2, are the target sites of BnaA03.WRKY28.

**Supplemental Figure 5.**
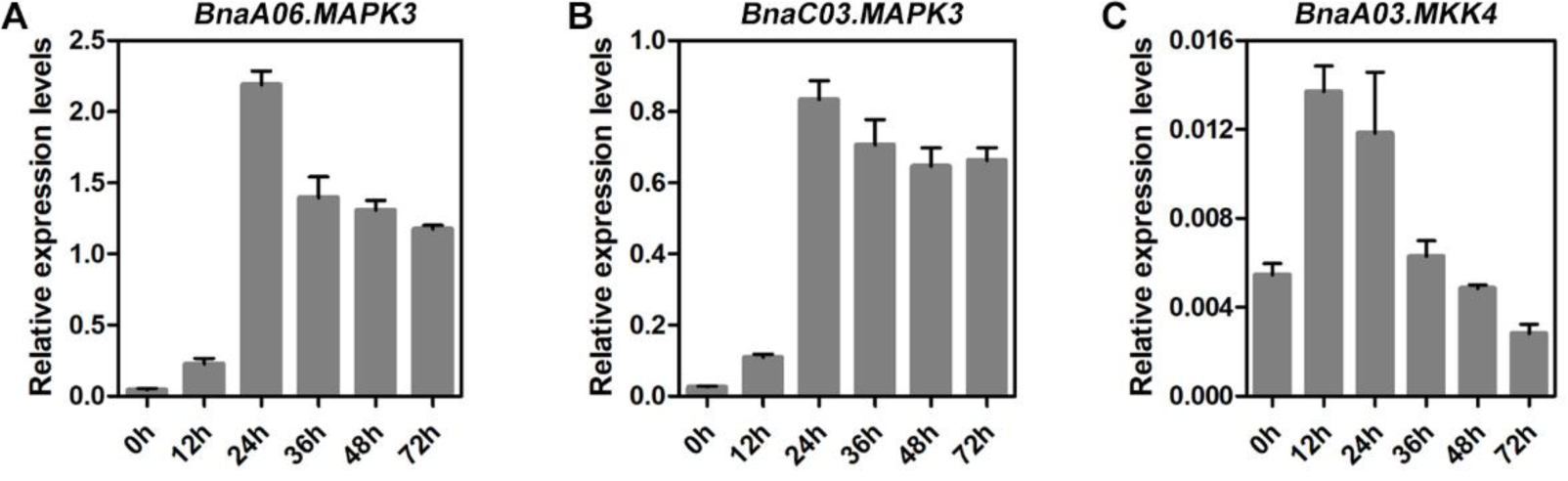
Expression profiling analysis of the MAPK cascade in response to *Sclerotinia* infection. (Supports Figure 4) **(A)** to **(C)** Induced expression profiles of *BnaA06.MPK3* **(A)**, *BnaC03.MPK3* **(B)**, and *BnaA03.MKK4* **(C)**, were identified at 0 h, 12 h, 24 h, 36 h, 48 h, and 72 h after inoculation with *S. sclerotiorum*. The latest fully unfolded leaves of rapeseed (Jia 9709) were used for infection in approximately 4 weeks. Data are shown as means ± SD (*n* = 3).

**Supplemental Figure 6.**
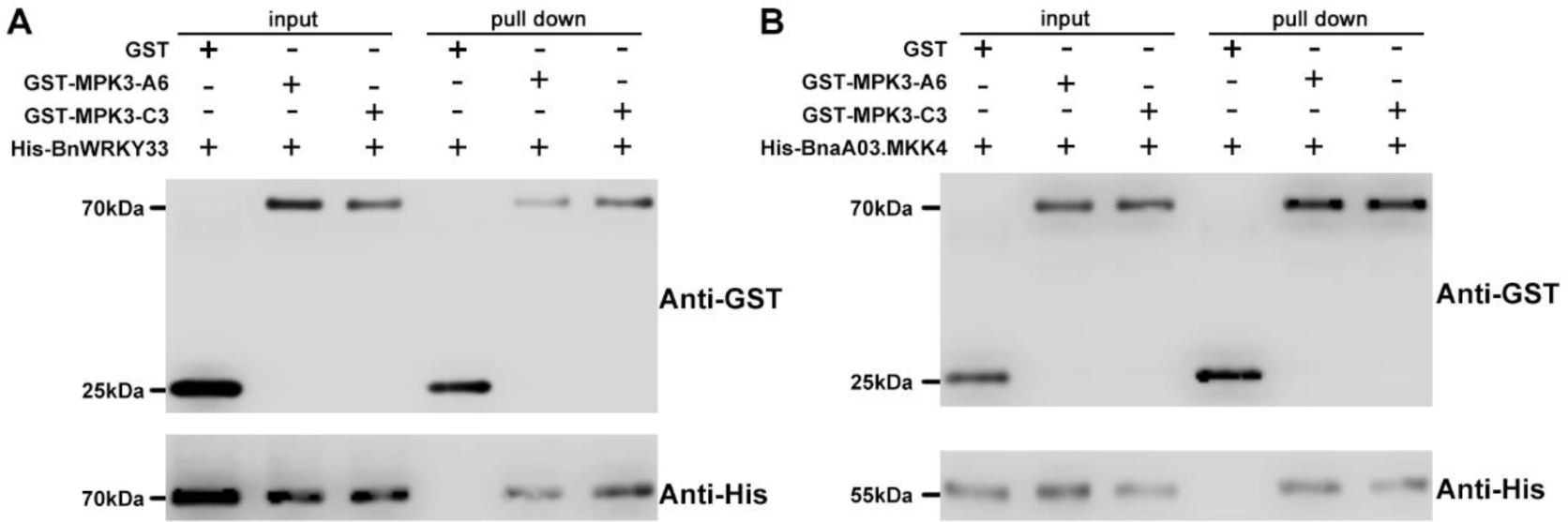
BnaA06.MPK3/BnaC03.MPK3 interacts with BnaA03.MKK4 and BnWRKY33 *in vitro*. (Supports Figure 4) **(A)** and **(B)** *In vitro* pull-down assays indicate that BnWRKY33 interacts with BnaA06.MPK3/BnaC03.MPK3 **(A)**, and BnaA03.MKK4 interacts with BnaA06.MPK3/BnaC03.MPK3 **(B)**, respectively. GST, GST-BnaA06.MPK3 (GST-MPK3-A6), or GST-BnaC03.MPK3 (GST-MPK3-A6) was immobilized on GST beads, incubated with His-BnWRKY33 or His-BnaA03.MKK4. The immunoblot assays were performed with anti-GST and anti-His antibodies.

**Supplemental Figure 7.**
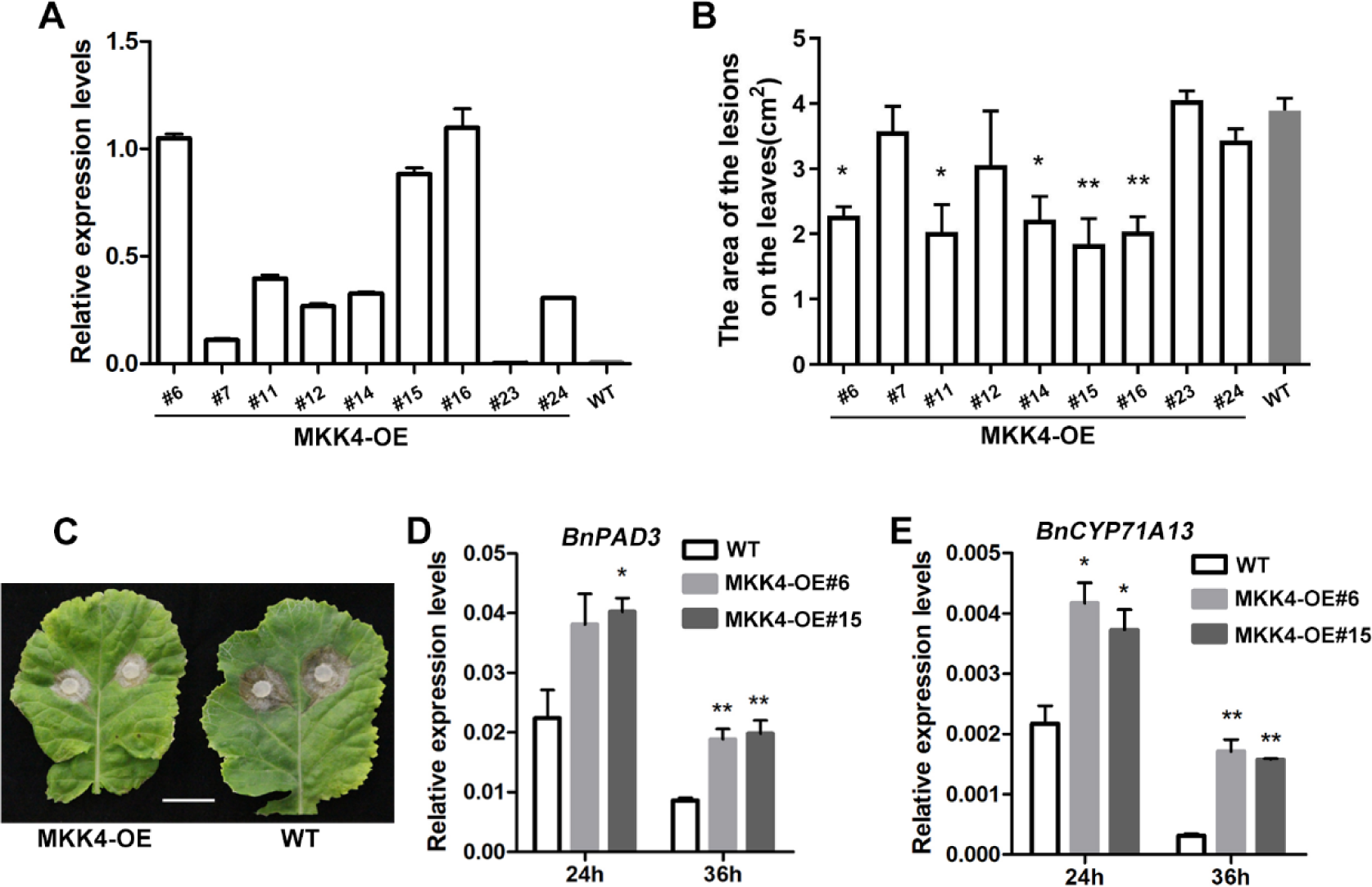
*BnaA03.MKK4* positively contributes to *S. sclerotiorum* resistance in *Brassica napus.* (Supports Figure 4 and Figure 5) **(A)** *BnaA03.MKK4* overexpression (MKK4-OE) lines establishment by qRT-PCR analysis. The expression in wild-type (WT) plants is the control. Data are shown as means ± SD (*n* = 3). **(B)** and **(C)** Statistical analysis of lesion areas of MKK4-OE lines after 48 h inoculation with *S. sclerotiorum*, and lesion imaging of representative lines. Scale bar, 2 cm. Data are shown as means ± SD (*n* = 3). Asterisks indicate significant differences compared with wild-type plants (t-test, \**P* < 0.05, \*\**P* < 0.01). **(D)** and **(E)** Relative expression levels of *BnPAD3*/*BnCYP71A13* in MKK4-OE lines and wild-type plants after *S. sclerotiorum* inoculation for 24 h or 36 h. Data are shown as means ± SD (*n* = 3). Asterisks indicate significant differences compared with wild-type plants at corresponding time points (t-test, \**P* < 0.05, \*\**P* < 0.01).

**Supplemental Figure 8.**
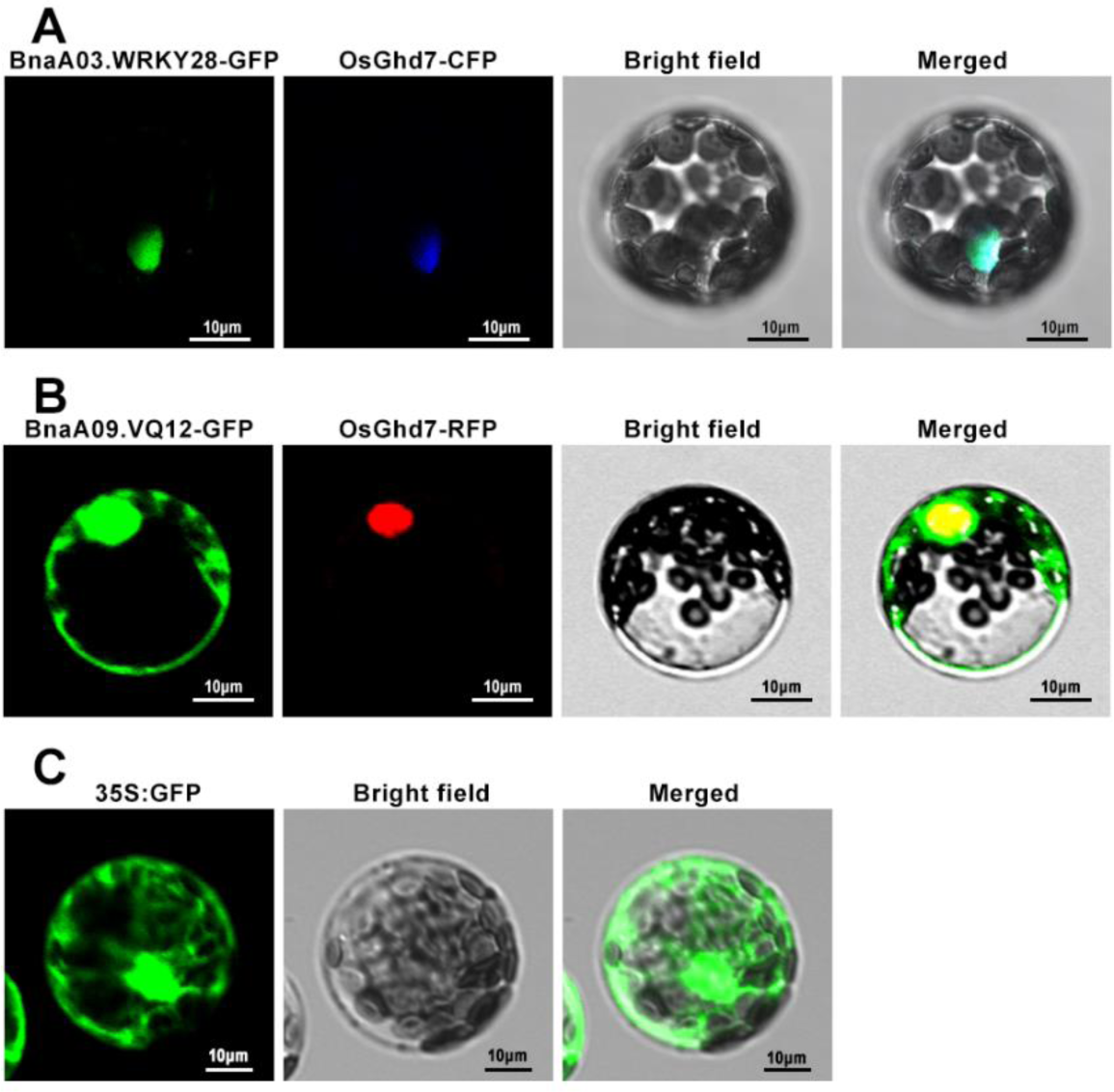
Subcellular localization of BnaA03.WRKY28 and BnaA09.VQ12 in *Arabidopsis* protoplasts. (Supports Figure 6) **(A)** Transcription factor BnaA03.WRKY28 was localized in the nucleus. OsGhd7-CFP was used as nuclear location signal. **(B)** BnaA09.VQ12 was localized in the cytoplasm and in the nucleus. OsGhd7- RFP was used as nuclear location signal. **(C)** The *Pro35S:GFP* was served as the positive control, which was distributed ubiquitously. In **(A)**, **(B)**, and **(C)**, scale bars, 10 μm.

**Supplemental Figure 9.**
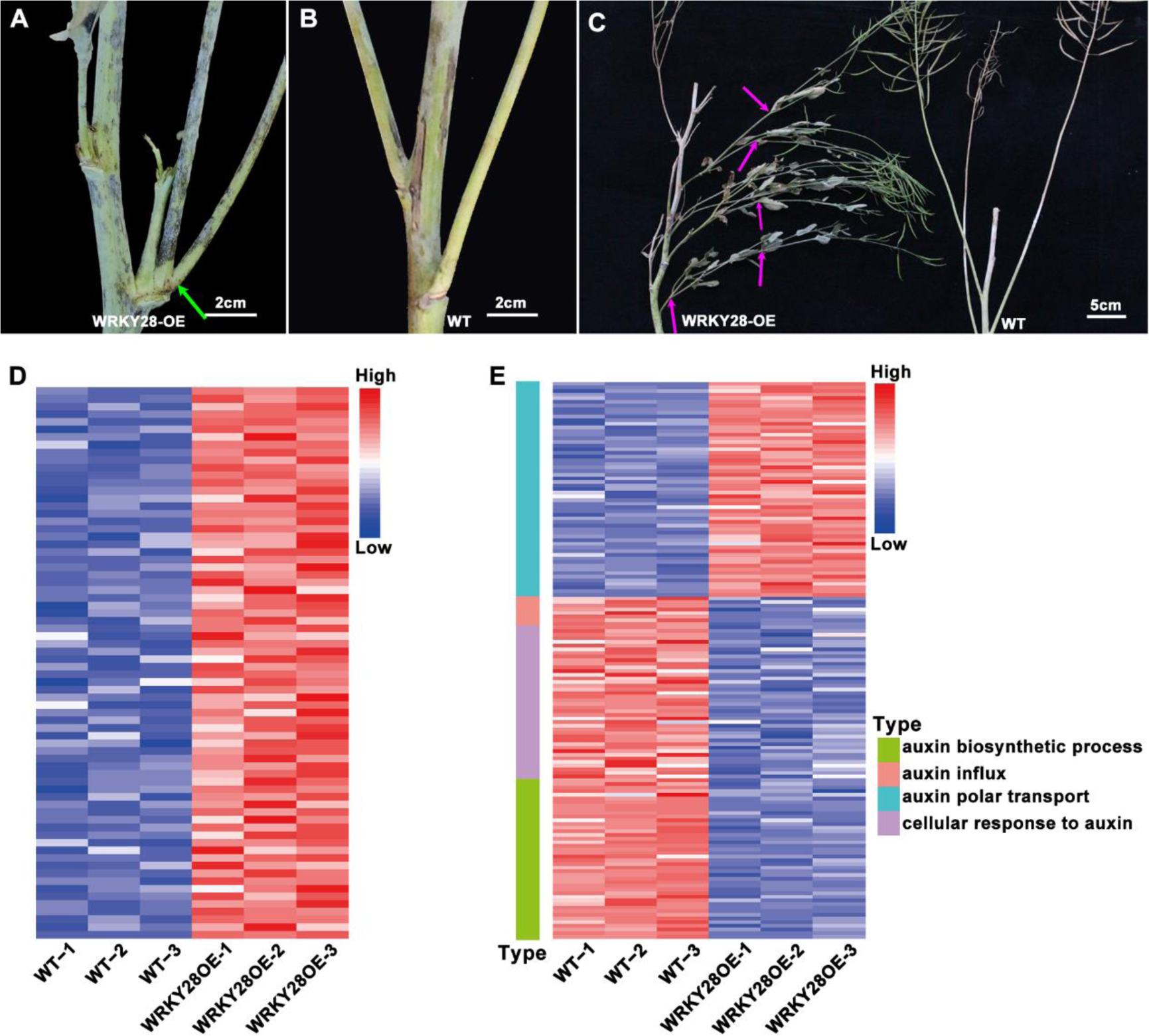
*BnaA03.WRKY28* overexpression lines exhibit excessive branches. (Supports Figure 9) **(A)** and **(B)** The formation of two branches in one leaf axil was obvious in *BnaA03.WRKY28* overexpression (WRKY28-OE) lines **(A)** compared with one branches in one leaf axil in the wild-type (WT) plants **(B)**. Scale bars, 2 cm. **(C)** The image shows when the rapeseed plants are severely damaged by *S. sclerotiorum*, the *BnaA03.WRKY28* overexpression lines produce more branches than the wild-type plants. Scale bar, 5 cm. **(D)** and **(E)** Heat maps showing the transcript levels of genes involved in meristem growth **(D)**, auxin biosynthetic process, auxin influx, auxin polar transport, and cellular response to auxin stimulus **(E)** in *BnaA03.WRKY28* overexpression lines and wild-type plants.

**Supplemental Figure 10.**
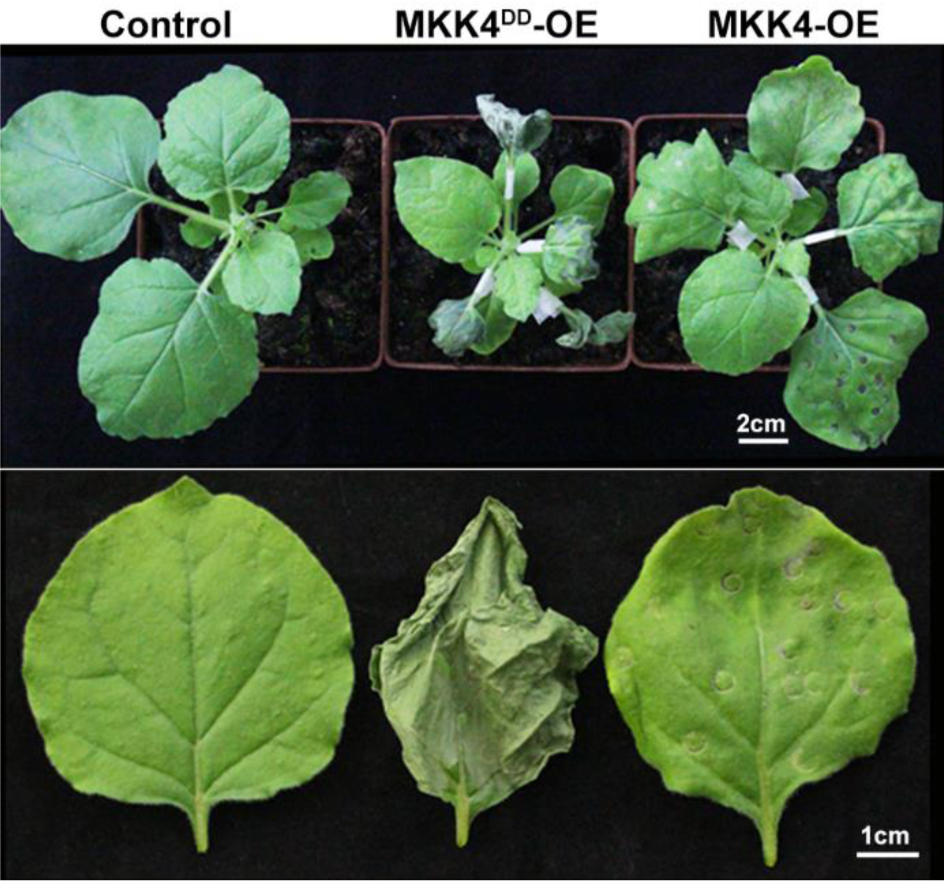
Constitutively active BnaA03.MKK4^DD^ induce hypersensitive response cell death. (Supports Figure 9F) Hypersensitive response cell death induced by overexpression of constitutively active BnaA03.MKK4^DD^ (MKK4^DD^-OE) in *N. benthamiana* leaves. MKK4-OE, plants infiltrated with *Agrobacterium* carrying *BnaA03.MKK4* overexpression driven by CaMV35S; MKK4^DD^-OE, plants infiltrated with *Agrobacterium* carrying *BnaA03.MKK4^DD^* overexpression driven by CaMV35S; Control, without injection. Scale bars, 5 cm (top image), 2 cm (bottom images).

